# TGFβ-induced long non-coding RNA *LINC00313* activates Wnt signalling and promotes cholangiocarcinoma

**DOI:** 10.1101/2022.09.28.509889

**Authors:** Panagiotis Papoutsoglou, Corentin Louis, Raphaël Pineau, Anaïs L’Haridon, Jesus M. Banales, David Gilot, Marc Aubry, Cédric Coulouarn

**Affiliations:** Inserm, Univ Rennes 1, OSS (Oncogenesis, Stress, Signalling) laboratory, UMR_S 1242, Centre de Lutte contre le Cancer Eugène Marquis, F-35042, Rennes, France; Department of Liver and Gastrointestinal Diseases, Biodonostia Health Research Institute, Donostia University Hospital, CIBERehd, Ikerbasque, San Sebastian, Spain; Department of Biochemistry and Genetics, School of Sciences, University of Navarra, Pamplona, Spain

**Keywords:** Cholangiocarcinoma, chromatin remodelling, LINC00313, lncRNA, TGFβ signalling, Wnt pathway

## Abstract

Cholangiocarcinoma (CCA) is a poor prognosis liver cancer characterized by high aggressiveness and resistance to therapy. Long non-coding RNAs (lncRNAs) and signals imposed by oncogenic pathways, such as transforming growth factor β (TGFβ), contribute to cholangiocarcinogenesis. Here, we identified LINC00313 lncRNA as a novel target of TGFβ signalling in CCA cells. TGFβ induced LINC00313 expression in a TβRI/Smad-dependent manner. Gene expression and genome-wide chromatin accessibility profiling revealed that nuclear *LINC00313* transcriptionally regulated genes involved in Wnt signalling, such as *TCF7*. *LINC00313* gain-of-function enhanced TCF/LEF-dependent transcription, promoted colony formation *in vitro* and accelerated tumour growth *in vivo*. Genes associated with *LINC00313* over-expression in human CCA were characterized by *KRAS* and *TP53* mutations and reduced patient’s overall survival. Mechanistically, actin-like 6A (ACTL6A), a subunit of SWI/SNF chromatin remodelling complex, interacted with *LINC00313* and impacted on *TCF7* and *SULF2* transcription. We propose a model whereby TGFβ induces *LINC00313* in order to regulate expression of hallmark Wnt pathway genes, in co-operation with SWI/SNF. By modulating key genes of the Wnt pathway, *LINC00313* fine-tunes Wnt/TCF/LEF-dependent transcriptional responses and boosts cholangiocarcinogenesis.

## Introduction

Cholangiocarcinoma (CCA) is an aggressive cancer from the biliary tree characterized by resistance to chemotherapies and poor prognosis. CCAs are categorized into intrahepatic (iCCA) and extrahepatic (eCCA), the later including perihilar (pCCA) and distal (dCCA) CCAs (Louis *et al*, 2020). CCA subtypes exhibit high tumour cell heterogeneity, as well as specific genetic and epigenetic alterations, including frequent *KRAS*, *TP53* and *IDH1/2* mutation and *FGFR2* gene fusions with enhanced tumourigenic activity (Kendall *et al*, 2019). Accordingly, signals imposed by oncogenic pathways, such as fibroblast growth factor (FGF), phosphatidylinositol-3-kinase (PI3K)-AKT and transforming growth factor β (TGFβ) frequently contribute to CCA development and/or progression (Fouassier *et al*, 2019).

TGFβ signals through type I and type II TGFβ receptors (TβRI and TβRII) that activate SMAD2, SMAD3 and SMAD4 which modulate gene expression (Tzavlaki & Moustakas, 2020). TGFβ-responsive targets include protein-coding and non-coding genes, which participate in a wide range of processes, such as liver fibrosis, tumour progression and metastasis (Papoutsoglou & Moustakas, 2020). TGFβ exerts both anti- and pro-tumourigenic functions. In normal epithelium, TGFβ restricts tumour initiation by promoting cell cycle arrest and apoptosis. However, in advanced malignancies, cancer cells become unresponsive to its tumour-suppressive actions, but still obey to its tumour-promoting functions, such as epithelial-to-mesenchymal transition (EMT) (Tu *et al*, 2019). In addition, TGFβ in the stroma facilitates cancer cell invasion and induces tumour immune evasion (Batlle & Massagué, 2019). In CCA, *SMAD4* is subjected to mutations, leading to gene inactivation. Loss of *SMAD4* is capable of nullifying the tumour suppressive axis of TGFβ. In addition, TGFβ is able to induce EMT and favour a fibrotic tumour microenvironment, thereby promoting metastasis (Papoutsoglou *et al*, 2019a). Notably, enhanced secretion of TGFβ2 in the stroma of iCCAs correlates with poor patient’s prognosis (Sulpice *et al*, 2013).

Long non-coding RNAs (lncRNAs) are transcripts longer than 200 nucleotides that lack protein-coding capacities. Long intergenic non-coding RNAs (lincRNAs) do not overlap with protein-coding genes. They form autonomous transcriptional units under the control of their own promoters. Although they structurally resemble messenger RNAs (mRNAs), functional open reading frames (ORFs) are not inherent to lncRNAs, rendering them incapable of encoding polypeptides. However, lncRNAs play crucial roles in physiological and pathological processes, including cancer. They do so by regulating molecular processes, such as gene transcription, splicing and translation, via interactions with DNA, proteins, mRNAs or microRNAs (miRNAs) (Yao *et al*, 2019). Several lncRNAs promote the proliferative, migratory and invasive properties of CCA cells (Jiang & Ling, 2019). Notably, lncRNAs with oncogenic functions, such as *lncRNA-ATB* (Lin *et al*, 2019), *CCAT1* (Zhang *et al*, 2017) and *TLINC* (Merdrignac *et al*, 2018) are over-expressed in CCA. Here, we focus on the role and the clinical relevance of *LINC00313* in CCA.

## Results

### *LINC00313* is a novel TGFβ target in CCA

By gene expression profiling, we identified 103 non-redundant genes differentially expressed by TGFβ in both HuCCT1 and Huh28 CCA cell lines (Fig. 1A), including known (*SERPINE1*) and novel coding and non-coding TGFβ targets (Table S1). Notably, the *long intergenic non-protein coding RNA 313* (*LINC00313*) was greatly induced by TGFβ in HuCCT1, Huh28 and NHC (Fig. 1A, B). *LINC00313* was up-regulated by TGFβ in a dose-dependent manner (Fig. 1C) and appeared to be an intermediate-to-late and early responsive gene in HuCCT1 and Huh28 cells, respectively (Fig. 1D). Using exon-specific primers, we further confirmed the induction of *LINC00313* by TGFβ in CCA cells (Fig. S1A). *LINC00313* was also highly expressed in hepatocellular carcinoma (HCC) but TGFβ had no impact on its expression in HepG2 and Hep3B HCC cell lines (Fig. S1B). Finally, subcellular fractionation demonstrated that *LINC00313* was mainly a nuclear lncRNA, similar to *MALAT1* and *SNORD48* (Fig. 1E).

**Fig. 1.**
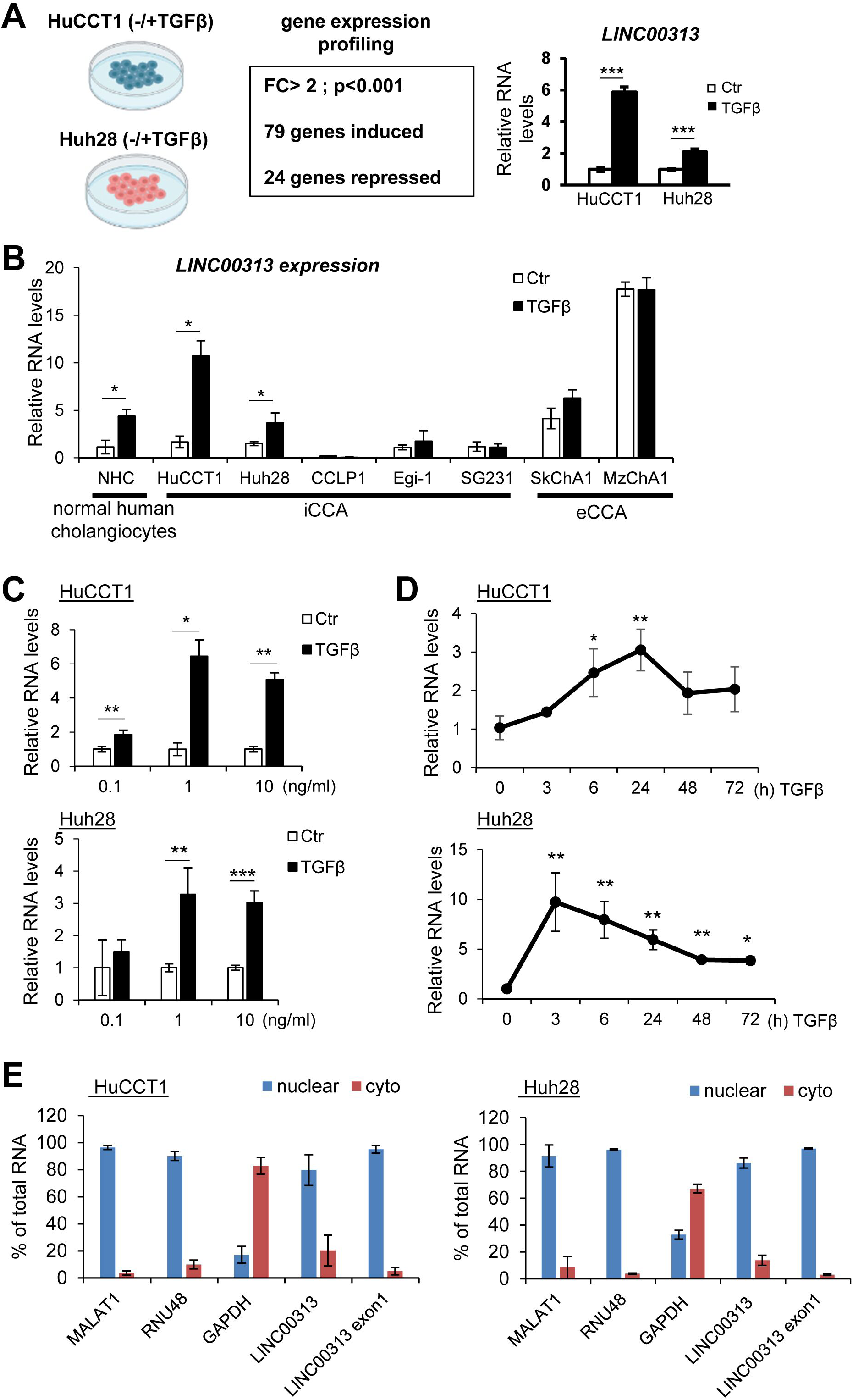
*LINC00313* is a TGFβ target gene. A) Experimental design to dissect TGFβ-regulated genes in CCA cells and *LINC00313* expression in HuCCT1 and Huh28 in response to TGFβ1, as identified by gene expression profiling. B) *LINC00313* expression in response to TGFβ1 stimulation for 16h, in NHC and CCA cell lines. C) *LINC00313* expression in response to the indicated doses of TGFβ1 in HuCCT1 and Huh28. D) *LINC00313* expression in response to TGFβ1 for the indicated time periods in HuCCT1 and Huh28. E) *MALAT1*, *RNU48*, *GAPDH* and *LINC00313* RNA levels in nuclear and cytoplasmic fractions of HuCCT1 and Huh28.

### TGFβ-induced *LINC00313* expression requires TβRI/Smad- and p38-dependent pathways

TGFβ signals through Smad-dependent (canonical) or Smad-independent (non-canonical) pathways (Tzavlaki & Moustakas, 2020). TβRI inhibitor LY2157299 abolished TGFβ-induced *LINC00313* expression in TGFβ-responsive cells (Fig. 2A, S1A, S2A). Similarly, blocking SMAD3 activation using SIS3 inhibitor completely abolished the TGFβ-mediated up-regulation of *LINC00313* in HuCCT1 and Huh28 cells (Fig. 2B). In HuCCT1, SMAD4 silencing reduced baseline and TGFβ-induced *LINC00313* expression similar to the combined SMAD2/3/4 silencing (Fig. 2C, S2B). In Huh28, SMAD2 silencing also prevented TGFβ-induced *LINC00313* expression (Fig. 2D, S2B).

**Fig. 2.**
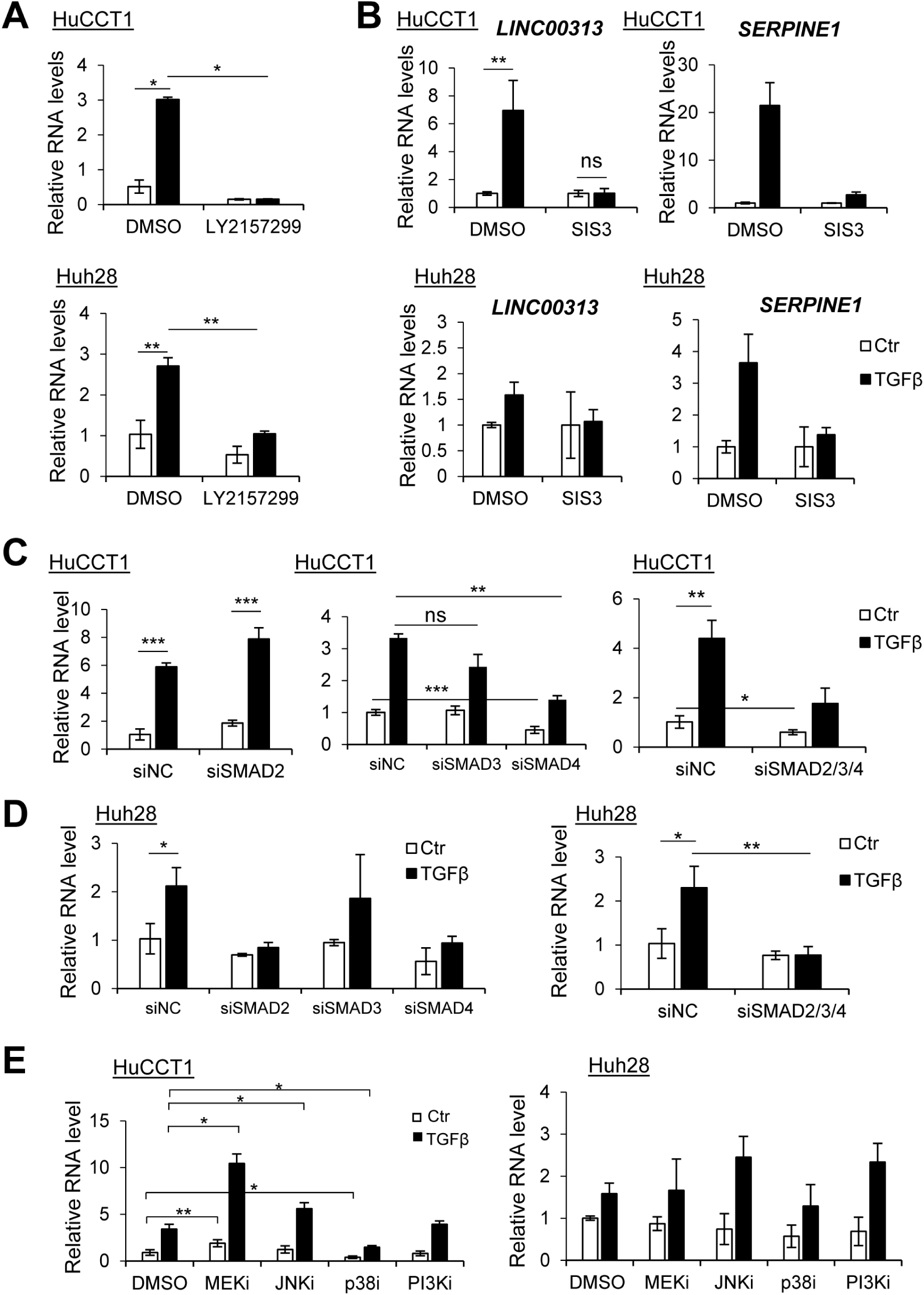
TGFβ induces *LINC00313* through TβRI/Smad-dependent and p38 kinase pathways. A) *LINC00313* expression in HuCCT1 and Huh28 cells treated with LY2157299 or DMSO, and stimulated with TGFβ1 or BSA/HCl. B) *LINC00313* and *SERPINE1* expression in HuCCT1 and Huh28 treated with SIS3 or DMSO and stimulated with TGFβ1 or BSA/HCl. C) *LINC00313* expression in HuCCT1 transiently transfected with siRNA targeting *SMAD2*, *SMAD3*, or *SMAD4*, alone or in combination, with or without TGFβ1. D) *LINC00313* expression in Huh28 transiently transfected with siRNA targeting *SMAD2*, *SMAD3*, or *SMAD4*, alone or in combination, with or without TGFβ1. E) *LINC00313* levels in HuCCT1 or Huh28 treated with MEK, p38, JNK or PI3K inhibitors, with or without TGFβ1 stimulation.

As regard to the non-canonical TGFβ pathway, p38 inhibition decreased, while MEK inhibition increased *LINC00313* expression. Inhibition of c-Jun N-terminal kinase (JNK) induced *LINC00313* in the presence of TGFβ in HuCCT1 cells (Fig. 2E). The efficiency of MAPKs inhibition was verified (Fig. S3A). The enhanced *LINC00313* expression upon MEK inhibition correlated with an enhanced TGFβ/SMAD response, as revealed by a SMAD-binding element (SBE) reporter assay (Fig. S3B). In Huh28 cells, *LINC00313* remained unchanged after blocking MAPKs (Fig. 2E). Thus, both SMAD3 and SMAD4 and the kinase activity of p38 are required for the TGFβ-mediated up-regulation of *LINC00313*.

### *LINC00313* modulates expression of genes involved in Wnt pathway

Based on its nuclear localization, we hypothesized that *LINC00313* acts at the transcriptional level to regulate gene expression (Fig. 3A). Gain-of-function tools were developed in HuCCT1 cells. An efficient *LINC00313* over-expression and a nuclear abundance similarly to parental HuCCT1 cells was confirmed (Fig. S4A). RNA-seq analysis identified 334 up- and 316 down-regulated genes after *LINC00313* gain-of-function (Fig. 3B, S4B, Table S2). Up-regulated genes were related to signalling pathways regulating pluripotency of stem cells (e.g. *WNT5A*, *TCF7*, *AXIN2*, *FZD2* and *ID1*) (Fig. 3C). Gene Set Enrichment Analysis (GSEA) further highlighted signatures of Hippo, Wnt and TGFβ pathways in the gene expression profile of *LINC00313* overexpressing cells (Fig. 3D, S4C). Induction of *WNT5A*, *AXIN2* and *SULF2* following *LINC00313* gain-of-function was also validated (Fig. 3E, S4D).

**Fig. 3.**
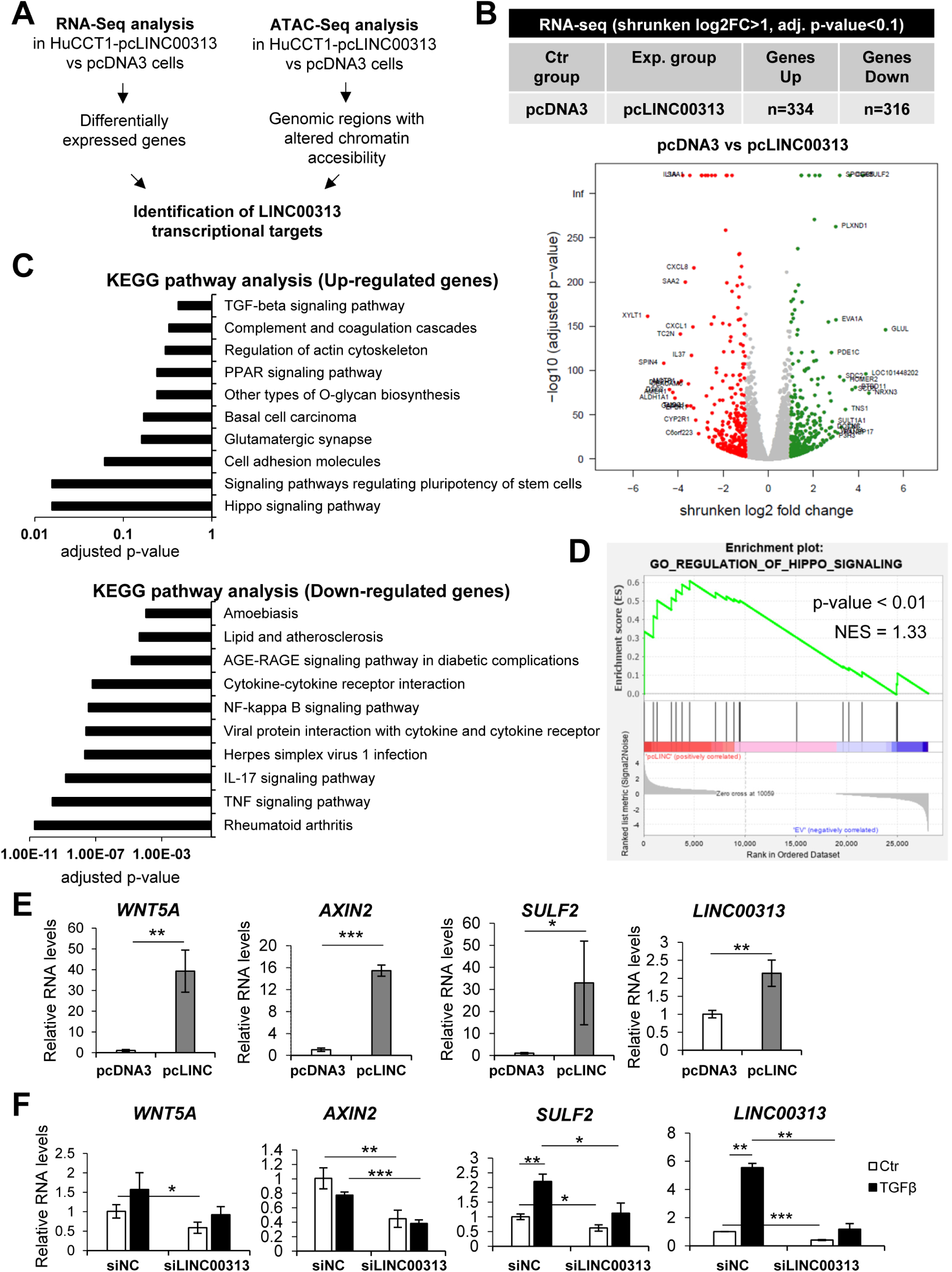
*LINC00313* regulates genes involved in the Wnt pathway. A) Experimental approach to identify *LINC00313* transcriptional targets. B) Number of differentially expressed genes in pcLINC00313 *versus* pcDNA3 HuCCT1 cells. Volcano plot shows differentially expressed genes. C) KEGG pathway analysis of up- or down-regulated genes. D) GSEA analysis of differentially expressed genes. E) *WNT5A*, *AXIN2* and *SULF2* mRNA levels in HuCCT1 expressing *LINC00313* or empty vector. F) *WNT5A*, *AXIN2* and *SULF2* expression in HuCCT1 transfected with siLINC00313 and stimulated with TGFβ1 or BSA/HCl.

Next, gene expression profiling after *LINC00313* silencing was performed in the presence or absence of TGFβ (Fig. S5A, S5B and Table S3). GO analysis confirmed that TGFβ-induced genes participated in cell migration, extracellular matrix organization and cell differentiation (Fig. S5C). KEGG pathway analysis revealed that many genes repressed after *LINC00313* silencing are associated with cancer signalling pathways regulating pluripotency of stem cells, including the Wnt signalling (Fig. S5D). In agreement with *LINC00313* gain-of-function experiments, *WNT5A*, *AXIN2* and *SULF2* expression levels were decreased upon *LINC00313* silencing in HuCCT1 cells (Fig. 3F). Expression of *TCF7*, *SULF2* and *WNT5A* was also diminished in *LINC00313*-silenced Huh28 and TFK1 cells (Fig. S6A). These observations were confirmed in Huh28 using another siRNA in order to exclude possible siRNA off-target effects (Fig. S6B).

### Nuclear *LINC00313* alters chromatin accessibility at genomic loci of Wnt-related genes

ATAC-seq identified genome-wide alterations in chromatin accessibility, including 23,628 unique peaks in *LINC00313* over-expressing cells (pcLINC00313) (Fig. 4A, S7A, S7B). No global alteration of the location of peaks, relative to genomic annotations was observed (Fig. S7C). Differential region analysis revealed 1657 genomic regions with increased accessibility and 2090 regions with decreased accessibility in pcLINC00313 cells (Fig. 4B and Table S4). HOMER motif enrichment analysis highlighted Fos-related antigen 1 (Fra1), an AP-1 transcription factor subunit, as the most significant enriched motif (Fig. S7D).

**Fig. 4.**
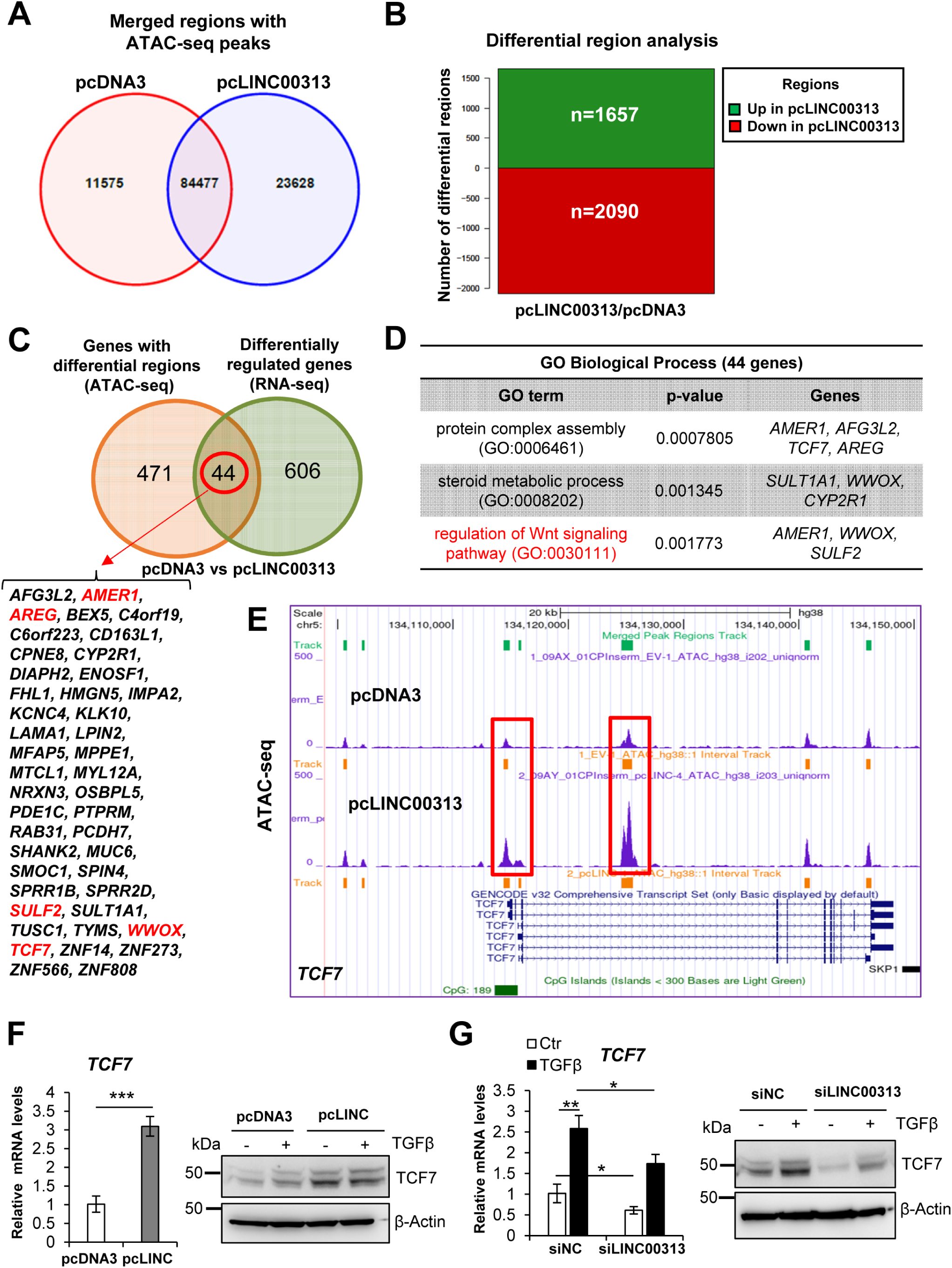
*TCF7* is transcriptionally induced by *LINC00313*. A) Venn diagram showing merged regions with overlapping and unique ATAC-seq peaks between pcDNA3 and pcLINC00313 expressing HuCCT1 cells. B) Bar plot illustrating the top differential regions. C) Venn diagram highlighting the 44 genes, whose chromatin accessibility and expression are altered. D) GO analysis of the 44 genes highlighted in panel C using Enrichr. E) Snapshot of the UCSC genome browser with ATAC-seq peaks at human *TCF7* gene locus, in pcDNA3 *versus* pcLINC00313 over-expressing HuCCT1 cells. F) *TCF7* mRNA and protein levels in HuCCT1 cells over-expressing LINC00313 or pcDNA3. G) *TCF7* RNA and protein levels in HuCCT1 cells, transfected with siLINC00313 and stimulated with TGFβ1.

We then integrated ATAC-seq and RNA-seq data and pinpointed the common genes. We found 44 genes that show both altered chromatin accessibility and altered expression after *LINC00313* over-expression (Fig. 4C). Interestingly, these genes were associated with the Wnt signalling pathway (Fig. 4D), known to play a key role in CCA progression(Boulter *et al*, 2015). Thus, we decided to investigate in deep the regulation of selected genes of the pathway, such as *TCF7*. Notably, we observed increased peak signal around the transcription start site and in the gene body of *TCF7* locus (Fig. 4E), as well as elevated *TCF7* mRNA and protein levels in *LINC00313* over-expressing cells (Fig. 4F). Interestingly, TGFβ induced *TCF7* expression whereas silencing *LINC00313* reduced *TCF7* mRNA and protein levels (Fig. 4G) in both control and TGF-β stimulated cells, suggesting that *LINC0313* may promote chromatin opening and enhanced transcriptional activity at the *TCF7* locus.

### *LINC00313* potentiates TCF/LEF transcriptional responses

Then, we investigated whether *LINC00313* could modulate Wnt/β-catenin-dependent transcription. We first validated luciferase reporter assays in HuCCT1 cells treated with CHIR99021 (CHIR), a glycogen synthase kinase 3 inhibitor, which activates the Wnt signalling. In HuCCT1 cells stably expressing a TCF/LEF reporter, CHIR increased reporter activity but TGFβ had no impact (Fig. S8A). Silencing *LINC00313* repressed CHIR-induced TCF/LEF reporter activity (Fig. 5A) and the associated target genes (Fig. 5B). *LINC00313* over-expression further enhanced the baseline and CHIR-induced TCF/LEF-luciferase activity (Fig. 5C). Administration of CHIR increased the viability of control HuCCT1 cells, an effect that was more pronounced in cells over-expressing *LINC00313* (Fig. S8B). Moreover, CHIR boosted colony formation of HuCCT1 control cells and to a larger magnitude of *LINC00313* over-expressing cells (Fig. S8C).

**Fig. 5.**
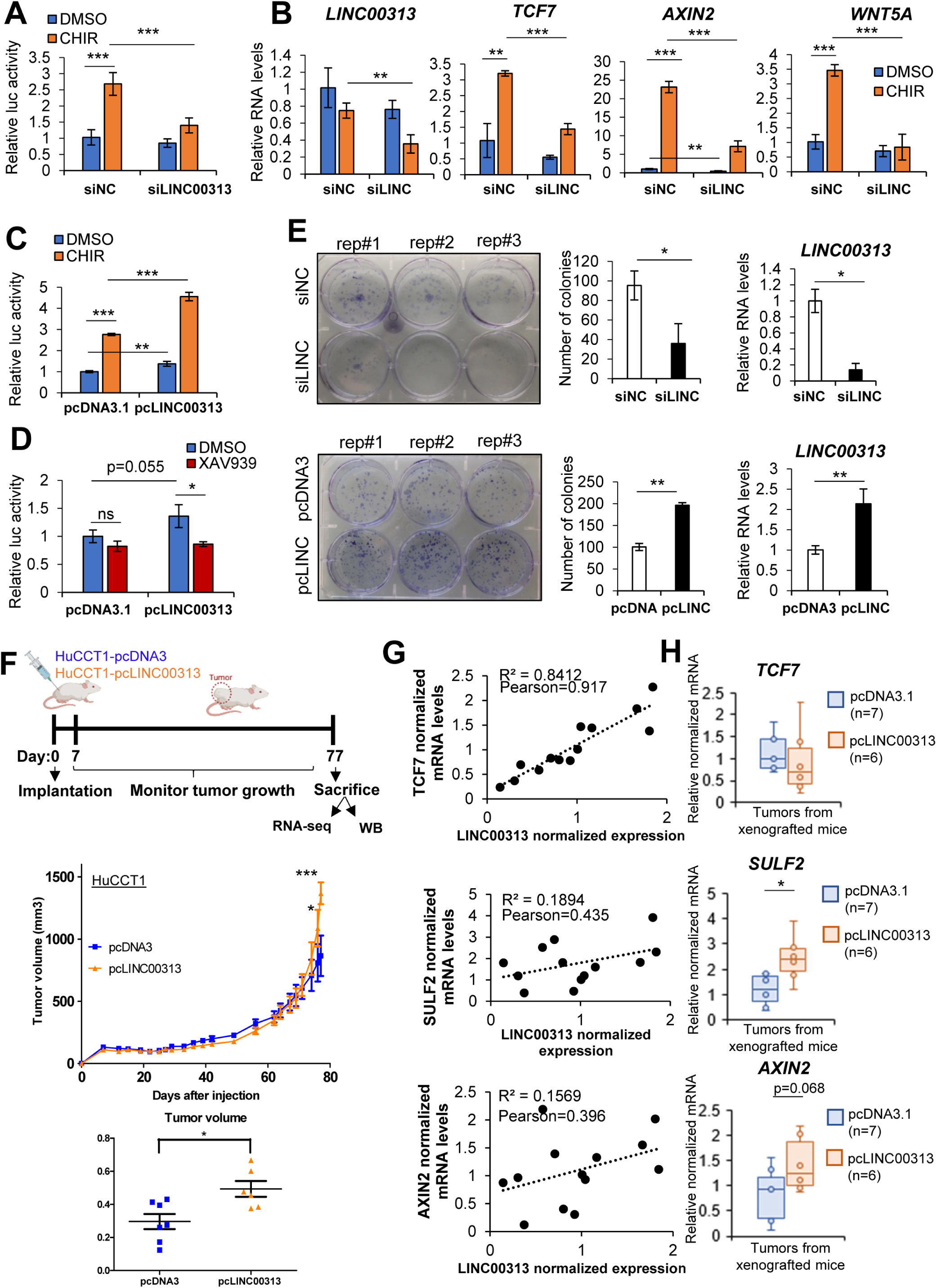
*LINC00313* modulates TCF/LEF-dependent transcriptional responses and enhances colony formation. A-B) TCF/LEF luciferase reporter assay (A) and qPCR analysis of *LINC00313*, *TCF7*, *AXIN2* and *WNT5A* expression (B) in HuCCT1 transiently transfected with siLINC00313 or siNC and treated with CHIR99021 or DMSO for 24h. C-D) TCF/LEF luciferase reporter assay in HuCCT1 stably over-expressing *LINC00313* or pcDNA3 and treated with CHIR99021 (C), XAV939 (D) or DMSO for 24h. E) Colony formation assay in HuCCT1 cells silenced or over-expressing *LINC00313*. F) *In vivo* mouse xenograft experiment, tumour growth curves and dot plot depicting the tumour volume in pcDNA3 (n=7) and pcLINC00313 (n=6) xenografts. G) Scatter plots depicting the correlation between *LINC00313* expression and each one of *TCF7*, *SULF2* and *AXIN2* mRNAs in RNA samples extracted from pcDNA3 and pcLINC00313 xenograft resected tumours. H) Boxplots of *TCF7*, *SULF2* and *AXIN2* expression.

XAV939, a Wnt pathway inhibitor, did not influence TCF/LEF responses in control cells, but reduced it in pcLINC00313 cells, which exhibit enhanced Wnt activity (Fig. 5D). Interestingly, XAV939 reduced colony formation (Fig. S8C). We confirmed the efficiency of the inhibitors by measuring *AXIN2* expression, as a typical Wnt-target gene. Indeed, CHIR strongly induced *AXIN2*, an effect that was enhanced in *LINC00313* over-expressing cells, whereas XAV939 modestly reduced *AXIN2* (Fig. S8D). Also, CHIR potentiated *TCF7* but not *LINC00313*, while XAV939 had no effect on their expression (Fig. S8D). β-catenin nuclear import is the major event that drives Wnt transcriptional responses. *LINC00313* over-expression did not change the subcellular localization of β-catenin (Fig. S8E). Collectively, the data support a model whereby *LINC00313* acts as a positive regulator of TCF/LEF-mediated signalling, but is dispensable for the initial steps of Wnt activation.

### *LINC00313* promotes CCA colony-forming capacities *in vitro* and tumour growth *in vivo*

Wnt signalling regulates cancer stem cell maintenance, cancer cell proliferation and migration (Rim *et al*, 2022). In this context, we showed that *LINC00313* boosted the ability of single cells to form colonies *in vitro* (Fig. 5E). Moreover, *LINC00313* over-expression neutralized the TGFβ-mediated decrease in HuCCT1 cell viability (Fig. S9A). However, *LINC00313* did not influence cell migration (Fig. S9B).

Interestingly, *LINC00313* accelerated tumour growth *in vivo* when cells were xenografted in nude mice (Fig. 5F, S10A). Expression analysis revealed a positive correlation between *LINC00313* and *TCF7*, *SULF2* and *AXIN2* mRNA levels in resected tumours (Fig. 5G). Moreover, we measured increased *SULF2* and *AXIN2* expression in *LINC00313* xenograft tumours, compared to control (Fig. 5H). Although *TCF7* mRNA levels did not change (Fig. 5H), we observed increased TCF7 protein expression in *LINC00313* tumours (Fig. S10B). *LINC00313* had no impact on the development of spontaneous metastases in mice (Fig. S10C) in agreement with the absence of effects on cell migration *in vitro*.

### The SWI/SNF complex subunit ACTL6A is an interactor of *LINC00313* lncRNA

RNA pull-down assay followed by mass spectrometry identified *LINC00313* partners possibly involved in chromatin remodelling. In total, 1538 proteins were identified to interact with *LINC00313* and 1840 proteins with the negative control *firefly luciferase* (*f-luc*) mRNA (Fig. 6A, Table S5). From the above, 121 proteins were specifically bound to *LINC00313* (Fig. 6B). In order to single out interesting candidates, we set three sorting layers (Fig. 6C). First, we focused on nuclear proteins (51/121), considering the nuclear localisation of *LINC00313*. Protein network analysis revealed multiple physical and functional associations and clustered them in four groups. The largest cluster contained proteins involved in cell cycle regulation and the rest members of the transcription factor TFIID complex, the integrator complex and proteins with RNA helicase activity (Fig. 6D). Second, we predicted the binding of *LINC00313* to 51 nuclear proteins using catRAPID (Agostini *et al*, 2013). Thirty two proteins were predicted to interact with *LINC00313* lncRNA (interaction propensity≥ 75, discriminative power≥ 98%) (Table S5). Third, among the 32 proteins we searched for transcription factors or chromatin modifiers (Fig. 6E), including actin-like 6A (ACTL6A) (Fig. 6F), a subunit of the SWI/SNF chromatin remodeling complex (Chang *et al*, 2021). Given that *LINC00313* modulates chromatin state of certain loci, we investigated the physical and functional association between *LINC00313* and ACTL6A deeper. Initially, we verified the nuclear localization of ACTL6A. TGFβ stimulation did not alter the nuclear abundance of ACTL6A (Fig. 6G). A specific interaction between *LINC00313* lncRNA and ACTL6A was demonstrated in HuCCT1, especially in the presence of TGFβ (Fig. 6H). Similarly, in HA-tagged ACTL6A over-expressing HEK293T cells, an even stronger interaction between *LINC00313* and HA-ACTL6A was observed (Fig. 6I). We also confirmed this interaction by RNA immunoprecipitation (RIP) assays. Interestingly, TGFβ stimulation increased enrichment of endogenous *LINC00313* to immunoprecipitated ACTL6A (Fig. 6J). Overall, *LINC00313* forms ribonucleoprotein complexes with several nuclear proteins that belong to transcriptional regulatory networks.

**Fig. 6.**
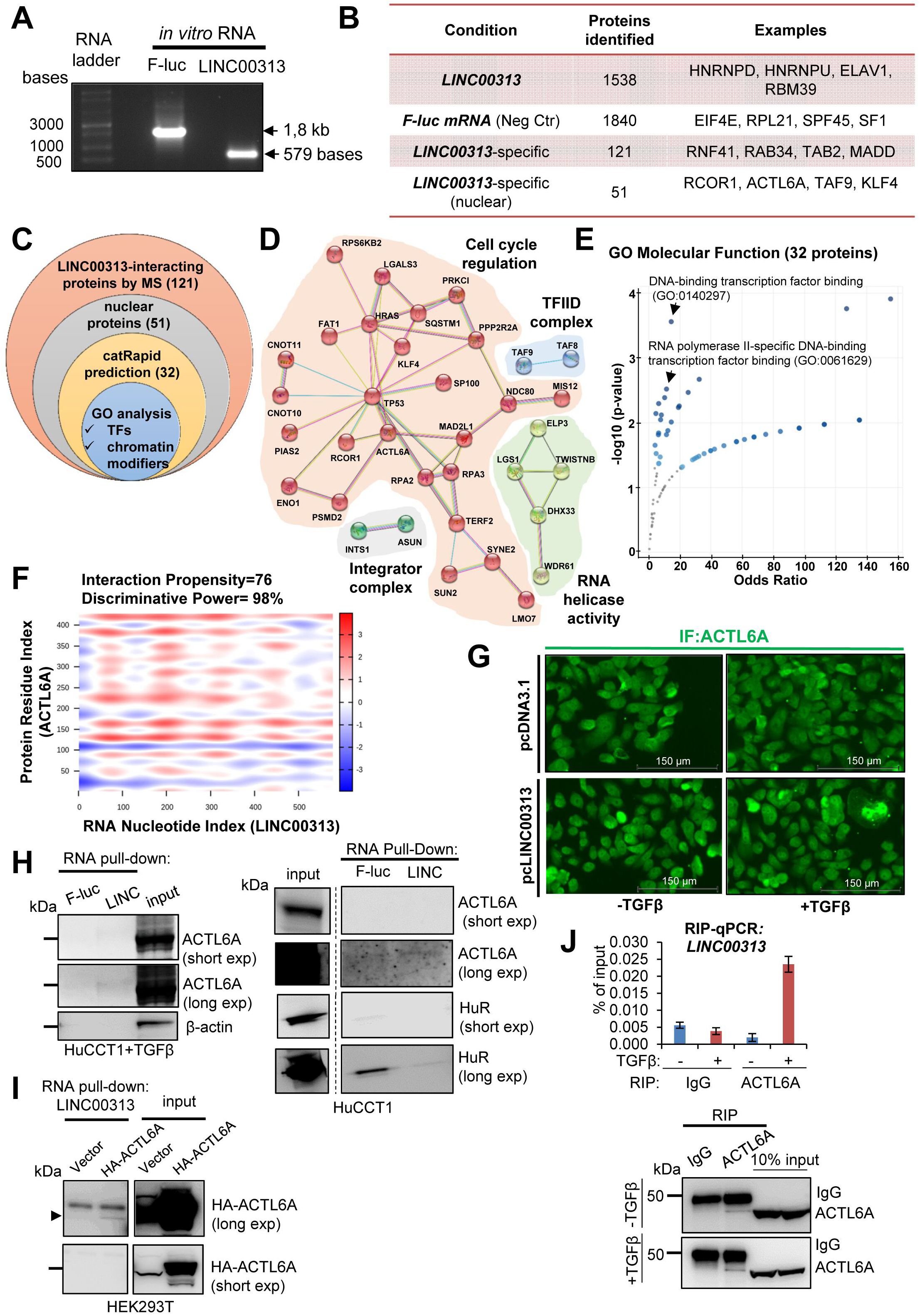
ACTL6A interacts with *LINC00313*. A) *In vitro* transcribed F-luc and *LINC00313* RNAs. B) Overview of interacting proteins identified by mass spectrometry. C) Venn diagram representing the strategy to narrow down *LINC0313*-binding proteins. D) Protein network analysis depicting physical and functional interactions between the 51 nuclear interactors. E) GO analysis of 32 nuclear interactors predicted by CatRAPID. F) Heatmap depicting the probability of interaction between *LINC00313* and ACTL6A. G) Immunofluorescence of ACTL6A in control or *LINC00313* over-expressing HuCCT1 cells, treated or not with TGFβ1. H) RNA pull-down assays in HuCCT1 cells, stimulated with TGFβ1 or not, using *in vitro* synthesized F-luc mRNA or *LINC00313* lncRNA, followed by immunoblotting for ACTL6A, HuR and β-actin. I) RNA pull-down assay in HEK293T cells, over-expressing an empty vector or HA-tagged ACTL6A using *in vitro* synthesized *LINC00313* lncRNA. J) RIP assay for endogenous ACTL6A followed by qPCR for *LINC00313* in HuCCT1 cells. An immunoblotting to verify the efficiency of ACTL6A immunoprecipitation is also shown.

### ACTL6A silencing or pharmacological inhibition of SWI/SNF complex diminishes TCF/LEF-mediated gene expression

Next, we evaluated the effects of ACTL6A on TCF/LEF-dependent transcription. Silencing ACTL6A attenuated *TCF7* mRNA and protein levels but did not affect TGFβ-induced *LINC00313* (Fig. 7A). ACTL6A silencing also resulted in decreased TCF/LEF luciferase activity and a drop in *TCF7* and *AXIN2* mRNA levels (Fig. 7B). Rescue experiments demonstrated that ACTL6A silencing dampened the *LINC00313*-mediated up-regulation of *SULF2* mRNA (Fig. 7C) and *TCF7* mRNA and protein levels (Fig. 7D).

**Fig. 7.**
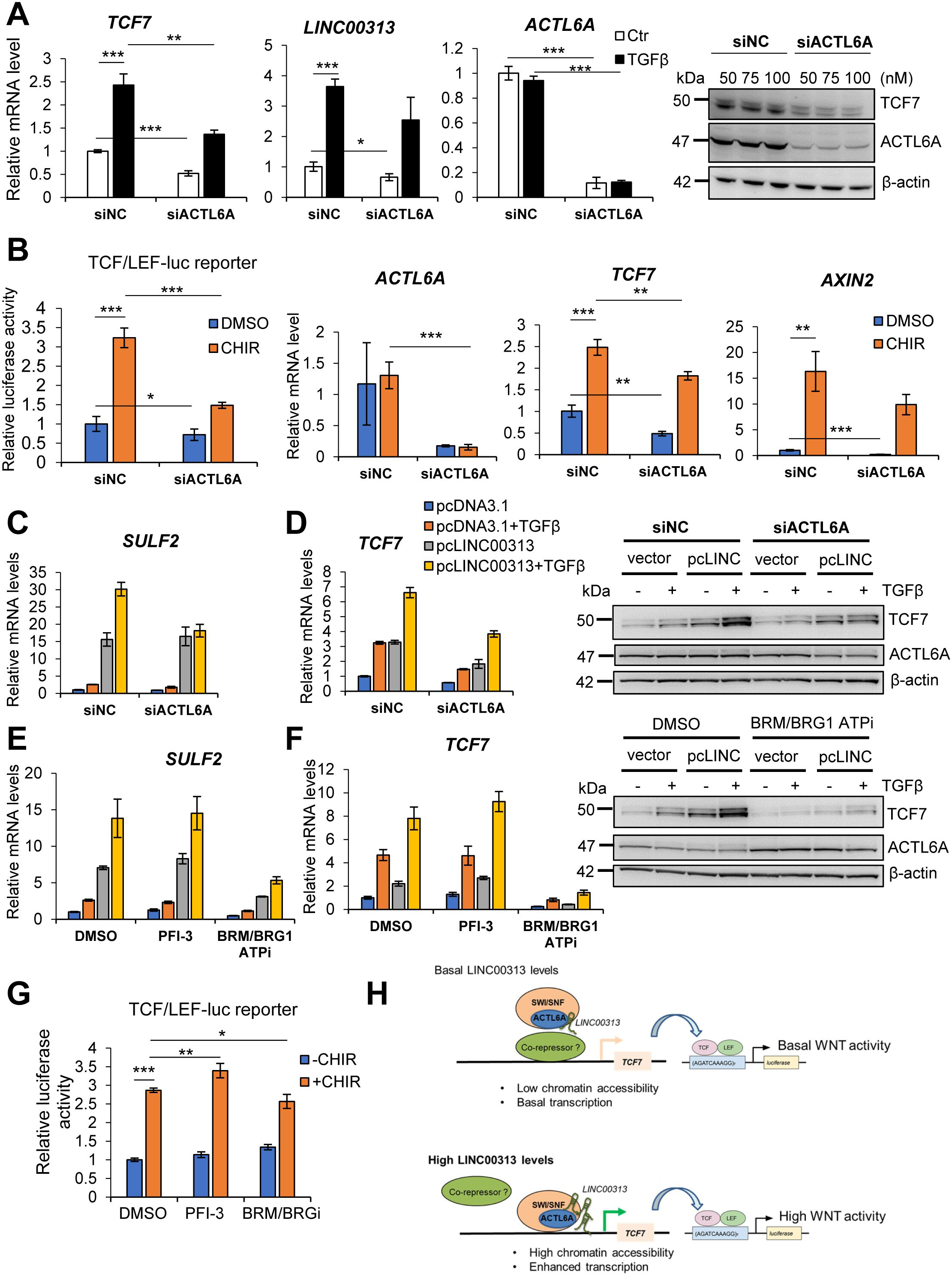
ACTL6A silencing or inhibition of SWI/SNF activity reduces TCF/LEF-mediated gene expression. A) *TCF7*, *LINC00313* and *ACTL6A* mRNA in HuCCT1 transfected with siACTL6A or siNC and in the presence or not of TGFβ1 and immunoblotting for TCF7, ACTL6A and β-actin. B) TCF/LEF luciferase reporter assay in siACTL6A or siNC HuCCT1 cells treated with CHIR99021 or DMSO for 24h and qPCR analysis of *ACTL6A*, *TCF7* and *AXIN2* mRNAs. C) *SULF2* mRNA in pcDNA3 or *LINC00313* over-expressing HuCCT1 transiently transfected with siACTL6A or siNC with or without TGFβ1. D) qPCR of *TCF7* mRNA and immunoblotting for TCF7, ACTL6A and β-actin in the same conditions as these of panel C. E) qPCR analysis of *SULF2* mRNA in pcDNA3 or *LINC00313* over-expressing HuCCT1 treated with PFI-3 or BRM/BRG ATPi or DMSO and with or without TGFβ1. F) qPCR analysis of *TCF7* mRNA and immunoblotting for TCF7, ACTL6A and β-actin in the same conditions as in panel E. G) TCF/LEF luciferase reporter assay in HuCCT1 cells treated with DMSO, PFI-3 or BRM/BRG ATPi and in the presence or not of CHIR99021. H) Proposed model for the molecular mechanism of *LINC00313*.

Since ACTL6A is an accessory subunit of SWI/SNF, we hypothesized that blocking the catalytic activity of the complex may yield effects similar to ACTL6A silencing. Thus, we utilized two inhibitors, targeting different domains of the core catalytic subunits SMARCA2/SMARCA4. Notably, the bromodomain inhibitor PFI-3 did not affect *SULF2* or *TCF7* mRNA expression. In contrast, administration of BRM/BRG1 ATPase inhibitor significantly reduced *SULF2* mRNA (Fig. 7E) and TCF7 expression both at the mRNA and protein levels (Fig. 7F), implying that these two genes are targets of SWI/SNF. Consistent with the effects on the individual genes, BRM/BRG1 ATPi, but not PFI-3 treatment resulted in a modest, but significant decrease of CHIR-induced TCF/LEF-luciferase reporter activity (Fig. 7G). Overall, we suggest a mechanism, whereby increased *LINC00313* levels promote chromatin accessibility in an ACTL6A/SWI/SNF-dependent manner and facilitate transcription of *TCF7* and *SULF2*, resulting in enhanced Wnt activation (Fig. 7H).

### *LINC00313* signature predicts poor prognosis in patients with CCA

From the TCGA dataset, *LINC00313* was neither overexpressed in CCA tissues nor correlated with overall or disease-free survival in patients with CCA (Fig. S11A). On the other hand, the expression levels of *ACTL6A*, *TCF7*, *WNT5A*, *AXIN2* and *SULF2* were all increased in CCA human tumours (Fig. S11B). Interestingly, *LINC00313* was significantly up-regulated in pancreatic ductal adenocarcinoma and correlated with lower disease-free survival (Fig. S11C).

Next, we hypothesized that if *LINC00313* expression may not be a prognostic factor in CCA, its activity could better reflect its clinical relevance. Thus, we performed RNA-seq analysis in control and *LINC00313* xenograft tumours to establish a signature reflecting *LINC00313* activity. In total, 347 genes were up-regulated and 327 genes were down-regulated in *LINC00313*-overexpressing tumours (Fig. 8A). Then, we integrated this signature with the gene expression profiles of 255 cases of clinically annotated human CCA. Integrative transcriptomics using principal component analysis (PCA) showed that the four control samples clustered together and were distinct to the four *LINC00313* samples (Fig. 8B). Dimension 2 of the PCA was related to *LINC00313* activity and identified two main clusters of human CCA (Fig. 8B). Interestingly, the cluster associated with *LINC00313* activity correlated with lower overall survival (Fig. 8C). Furthermore, *IDH1* and *IDH2* mutations were more prevalent in control cluster, while *KRAS* and *TP53* mutations were significantly enriched in *LINC00313* cluster (Fig. 8D). We also observed increased levels of carbohydrate antigen 19-9 and γ-glutamyltransferase and decreased levels of albumin, markers of hepatobiliary disease and of unfavourable prognosis in CCA, in the *LINC00313* cluster (Fig. 8E). In addition, the *LINC00313* cluster was characterized by significantly higher perineural invasion and regional lymph node metastasis (Fig. 8E). Collectively, we conclude that although *LINC00313* expression *per se* is not a prognostic factor in CCA, the *in vivo* gene signature that reflects its activity exhibits a strong prognostic value in terms of survival outcome.

**Fig. 8.**
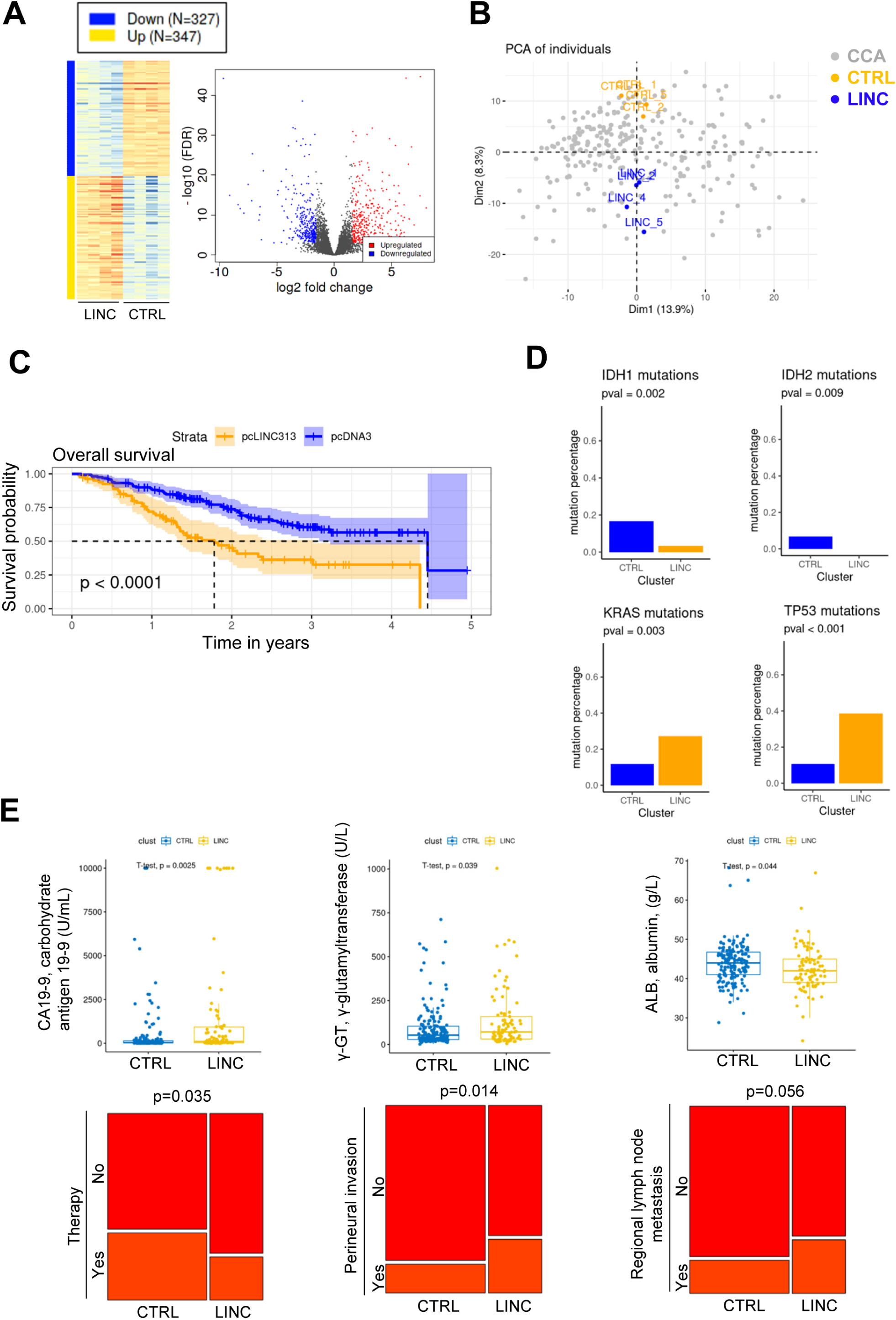
*In vivo LINC00313*-associated gene signature predicts survival outcome of CCA patients. A) Heatmap and volcano plot showing differentially expressed genes in control (n=4) and LINC00313 (n=4) resected xenograft tumours. B) PCA of control and LINC00313 samples as described in panel A and human CCA samples. C) Survival curve depicting overall survival of CCA patients in control and LINC00313 clusters. D) Mutational analysis showing the percentage of *IDH1*, *IDH2*, *KRAS* and *TP53* mutations. E) Clinical parameters of CCA patients in control and LINC00313 clusters.

## Discussion

By exploring the TGFβ-regulated transcriptome in CCA cell lines, we identified *LINC00313* as a TGFβ-responsive gene, a finding that expands the growing list of TGFβ-regulated lncRNAs (Papoutsoglou & Moustakas, 2020). An interesting observation is that TGFβ is able to induce *LINC00313* only in a subset of CCA cell lines, as well as in normal cholangiocytes. A possible explanation is that some CCA cell lines are not responsive to TGFβ stimulation. Since most of the CCA cell lines have not, yet, been fully characterized, concerning the responsiveness to TGFβ, a deeper investigation on TGFβ family members is required. On the other hand, TGFβ did not affect *LINC00313* expression in HepG2 and Hep3B, two HCC cell lines known to respond to TGFβ signalling, suggesting that TGFβ may selectively regulate this gene in certain cancer cells.

*LINC00313* is a lincRNA that possesses oncogenic properties and is a marker of poor prognosis in papillary thyroid cancer (Wu *et al*, 2018; Yan *et al*, 2019). Also, a pro-tumorigenic role of *LINC00313* has been described in osteosarcoma (Xing *et al*, 2022) and cervical carcinoma (Zhai *et al*, 2021). Moreover, *LINC00313* is up-regulated in glioma and promotes tumorigenesis, via enhancing cell proliferation, migration and invasion (Shao *et al*, 2019). The functional role of *LINC00313* in CCA remained unknown but our data support a pro-oncogenic role. We demonstrated that *LINC00313* favours CCA tumour growth *in vivo* and colony formation *in vitro*. Although we did not observe up-regulation of *LINC00313* in human CCA, we showed that *in vivo* gene signature associated with *LINC00313* gain-of-function correlated with poor overall survival. We speculate that *LINC00313* activity, rather than its expression, is of paramount importance for the stratification of CCA patients based on survival outcomes.

Integration of ATAC-seq and RNA-seq data allowed us to select genes involved in Wnt signalling. The Wnt pathway plays crucial roles in CCA progression and is often activated in CCA tissues. CCA tumours are usually characterized by enhanced expression of several Wnt ligands and increased nuclear accumulation of β-catenin (Selvaggi *et al*, 2022). We focused on *TCF7* because it encodes one of the main transcription factors of Wnt/β-catenin pathway (Doumpas *et al*, 2019). Furthermore, *TCF7* reinforces CCA progression by positively regulating MYC and FOSL1(Liu *et al*, 2019) and induces CCA proliferation and drug resistance, via regulating SOX9/FGF7/FGFR2 axis (Liu *et al*, 2022). *SULF2* is the second interesting transcriptional target of *LINC00313*. It encodes a heparan sulfate-editing enzyme that activates Wnt signalling, by facilitating the bioavailability of Wnt ligands to their receptors (Rosen & Lemjabbar-Alaoui, 2010). *SULF2* is up-regulated in human CCA and is associated with enhanced PDGFRβ-YAP signalling, tumour progression and chemoresistance. Importantly, targeting SULF2 protein with a monoclonal antibody abolishes tumour growth in a mouse CCA xenograft model (Luo *et al*, 2021). We also identified additional genes induced by *LINC00313*, such as *ID1* and *ID3*, which inhibit cell differentiation. Thus, we suggest that *LINC00313* positively regulates genes related to cell proliferation and stemness that could explain its effects on colony formation and CCA xenograft tumour growth.

Our proteomic analysis revealed numerous nuclear proteins that associate with *LINC00313*. We focused on ACTL6A, which is frequently over-expressed in cancer. For instance, ACTL6A is up-regulated in HCC and promotes migration and invasion *in vitro*, as well as tumour growth and metastasis *in vivo*, via activating Notch signalling (Xiao *et al*, 2016). The role of ACTL6A in CCA is still unknown. However, the SWI/SNF catalytic subunit BRG1 is over-expressed and predicts a poor prognosis in iCCA. More importantly, it has been demonstrated that BRG1 activates the Wnt pathway through transcriptional regulation of Wnt receptor and target genes, but also via binding to β-catenin/TCF4 transcription complex, in HuCCT1 (Zhou *et al*, 2021). These observations are in line with our data indicating decreased activation of Wnt transcriptional responses upon *ACTL6A* silencing or BRG1 inhibition.

At the molecular level, TGFβ promotes ribonucleoprotein complexes between *LINC00313* and ACTL6A without changing ACTL6A expression. This finding opens up the possibility that TGFβ may engage *LINC00313* to bring the chromatin remodelling machinery, in close proximity to Smads, at the regulatory regions of selected targets. Although ACTL6A is not a classical RNA-binding protein, it was previously reported to interact with the lncRNA uc.291 to regulate epidermal differentiation (Panatta *et al*, 2020).

Several lncRNAs establish interactions with SWI/SNF subunits. For example, SWINGN lncRNA binds to SMARCB1 and facilitates transcription of target genes (Grossi *et al*, 2020). In HCC cells, lncTCF7 interacts with BRG1, BAF170 and SNF5 subunits and recruits SWI/SNF to *TCF7* promoter, thereby activating *TCF7* transcription *in cis* (Wang *et al*, 2015). Our mechanistic evidence propose a model, whereby *LINC00313* acts *in trans* to regulate *TCF7* and *SULF2* transcription, through binding ACTL6A and possibly via increased loading of SWI/SNF to *TCF7* and *SULF2* loci, setting an additional layer of regulation of TCF/LEF signalling. Undoubtedly, additional modes of action for *LINC00313* cannot be excluded. For example, nuclear lncRNAs frequently form triplex formation with DNA at chromatin regions, through direct base pairing (Zapparoli *et al*, 2020). It is worth to investigate whether *LINC00313* participates in these loops, in order to alter chromatin accessibility. Nevertheless, our study highlights an important and new role for *LINC00313* as a modulator of Wnt/TCF signalling, that affects tumour progression and with clinical implications in CCA.

## Materials and Methods

### Cell culture

Primary cultures of normal human cholangiocytes (NHC) were established, characterized and cultured as previously described (Urribarri *et al*, 2014). HuCCT1 and Huh28 CCA cell lines were from RIKEN BioResource Center (Tsukuba-shi, Japan) and CCLP1, Egi-1, SG231, TFK1, Sk-ChA-1 and Mz-ChA-1 were provided by Laura Fouassier (Paris, France). HepG2/C3A and HEK293T were from ATCC (www.lgcstandards-atcc.org). Serum-starved cells were stimulated with human recombinant TGFβ1 (R&D Systems) for 16h.

### Treatments with inhibitors

The TβRI inhibitor LY2157299 (Interchim, LSK040) was added to the cells at a final concentration of 10 µM. The SMAD3 inhibitor (E)-SIS3 (MedChemExpress, HY-13013) was used at a final concentration of 10 µM. Cells were treated with 0.5 µM PD184352 (Sigma-Aldrich, PZ0181), a MEK inhibitor, 10 µM SP600125 (Sigma-Aldrich, S5567), a JNK inhibitor, 10 µM SB203580 (Sigma-Aldrich, S8307), a p38 inhibitor and 0.1 µM wortmannin (Sigma-Aldrich, W1628), a PI3K inhibitor. All treatments with the inhibitors were performed 1h prior the addition of TGFβ1. The GSK3 inhibitor CHIR-99021 (TargetMol, T2310) and the tankyrase inhibitor XAV-939 (TargetMol, T1878) were used at 10 µM final concentration. The SMARCA2/4 bromodomain inhibitor PFI-3 (MedChemExpress, HY-12409) and the allosteric dual SMARCA2 and brahma related gene 1 (BRG1)/SMARCA4 ATPase activity inhibitor BRM/BRG1 ATP Inhibitor-1 (MedChemExpress, HY-119374) were administrated to cells at 10 and 5 µM final concentrations, respectively. Dimethyl sulfoxide (DMSO) was used as a vehicle control treatment.

### Plasmid transfections

Human *LINC00313* (NR_026863.1) was synthesized and cloned into the pcDNA3.1/Hygro (+) expression vector (Invitrogen), by Eurofins (Eurofins Genomics, Germany). The sequence of the plasmid was verified by double strand DNA sequencing. For the generation of stable *LINC00313* over-expressing HuCCT1 clones, cells were transfected with pcDNA3.1-LINC00313 for 48 h. Then, transfected cells were grown, for 2 weeks, in fresh selection medium, consisting of 10% FBS/RPMI, in the presence of 650 µg/ml hygromycin B Gold (Invivogen, ant-hg-1). By using limiting dilution assay, individual clones from the stable LINC00313 over-expressing pool were seeded in 96-well plates and grown in selection medium. The same protocol was followed, in order to establish stable HuCCT1 clones over-expressing an empty vector (pcDNA3.1), which served as control for gain-of-function experiments. Human HA-tagged ACTL6A plasmid (VB211129-1064fwh) was constructed by Vectorbuilder. M50 Super 8x TOPFlash (Addgene plasmid #12456) and M51 Super 8x FOPFlash (TOPFlash mutant) (Addgene plasmid #12457) were a gift from Randall Moon. HuCCT1 or HEK293T cells were transfected with plasmid DNA, using Lipofectamine 2000 reagent (ThermoFisher Scientific, 11668019), according to the instructions by the manufacturer.

### Lentiviral infection

The infection of HuCCT1 cells with pGreenFire 2.0 TCF/LEF reporter virus (pGF2-TCF/LEF-rFluc-T2A-GFP-mPGK-Puro) (SBI System Biosciences, TR413VA-P) was performed using SureENTRY Transduction Reagent. Cells were infected at a final multiplicity of infection (MOI) of 10. The next day medium was replaced with fresh complete medium and cells were incubated for another 24 h. Two days after transduction, cells were treated with 1 μg/ml puromycin for two weeks for selection of infected cells.

### siRNA transfections

HuCCT1 cells were seeded at 70% confluency and transfected with siRNAs at a final concentration of 50 nM. For single transfections with two different siRNAs, each siRNA was used a concentration of 25 nM. Similarly, for single transfections with three different siRNAs simultaneously, each siRNA was used a concentration of 25 nM, so that the final siRNA concentration reached to 75 nM. Transient siRNA transfections were performed using Lipofectamine RNAiMAX transfection reagent (ThermoFisher Scientific, 13778075), according to the manufacturer’s instructions. Two custom-made siRNAs (Dharmacon, CTM-484556 and CTM-632606) were designed to target *LINC00313* (NR_026863.1). ON-TARGETplus human SMARTpool siRNAs were used for silencing *SMAD2*, *SMAD3*, *SMAD4* and *ACTL6A*. For negative control transfections (siNC), an ON-TARGETplus Non-targeting Pool was used. A complete list of the siRNAs used in this study is provided in supplementary Table S6.

### ATAC sequencing

The ATAC-seq experiment was performed using the ATAC-seq service from Active Motif. Briefly, 100,000 HuCCT1 pcDNA3.1 (empty vector) and pcLINC00313 over-expressing cells were used per ATAC reaction. Cells were harvested and frozen in culture media containing FBS and 5% DMSO. Cryopreserved cells were sent to Active Motif to perform the ATAC-seq assay. The cells were then thawed in a 37 °C water bath, pelleted, washed with cold PBS, and tagmented as previously described. Briefly, cell pellets were resuspended in lysis buffer, pelleted, and tagmented using the enzyme and buffer provided in the Nextera Library Prep Kit (Illumina). Tagmented DNA was then purified using the MinElute PCR purification kit (Qiagen), amplified with 10 cycles of PCR, and purified using Agencourt AMPure SPRI beads (Beckman Coulter). Resulting material was quantified using the KAPA Library Quantification Kit for Illumina platforms (KAPA Biosystems), and sequenced with PE42 sequencing on the NovaSeq 6000 sequencer (Illumina).

### RNA sequencing

Total RNA from HuCCT1 pcDNA3.1 (empty vector) and pcLINC00313 over-expressing clones was isolated using the miRNeasy kit (Qiagen, 217004). RNA-seq was performed by Active Motif. For each sample, 2 ug of total RNA was then used in Illumina’s TruSeq Stranded mRNA Library kit (Cat# 20020594). Libraries were sequenced on Illumina NextSeq 500 as paired-end 42-nt reads. Sequence reads were analyzed with the STAR alignment. The data analysis was performed as follows:

Read Mapping: The paired-end 42 bp sequencing reads (PE42) generated by Illumina sequencing (using NextSeq 500) are mapped to the genome using the STAR algorithm with default settings.

Fragment Assignment: The number of fragments overlapping predefined genomic features of interest (e.g. genes) are counted. Only read pairs that have both ends aligned are counted. Read pairs that have their two ends mapping to different chromosomes or mapping to same chromosome but on different strands are discarded. The gene annotations used were obtained from Subread package. These annotations were originally from NCBI RefSeq database and then adapted by merging overlapping exons from the same gene to form a set of disjoint exons for each gene. Genes with the same Entrez gene identifiers were also merged into one gene.

Differential Analysis: After obtaining the gene table containing the fragment counts of genes, we perform differential analyses to identify statistically significant differential genes using DESeq2. The following lists the pre-processing steps before differential calling. a. Data Normalization: DESeq2 expects un-normalized count matrix of sequencing fragments. The DESeq2 model internally corrects for library size using their median-ofratios method. The gene table obtained from Analysis Step 2) is used as input to perform the DESeq2’s differential test. b. Filtering before multiple testing adjustment: After a differential test has been applied to each gene except the ones with zero counts, the p-value of each gene is calculated and adjusted to control the number of false positives among all discoveries at a proper level. This procedure is known as multiple testing adjustment. During this process, DESeq2 by default filters out statistical tests (i.e. genes) that have low counts by a statistical technique called independent filtering. It uses the average counts of each gene (i.e. baseMean), across all samples, as its filter criterion, and it omits all genes with average normalized counts below a filtering threshold from multiple testing adjustment. This filtering threshold is automatically determined to maximize detection power (i.e. maximize the number of differential genes detected) at a specified false discovery rate (FDR).

### cDNA synthesis and real-time qPCR

Total RNA was isolated using the miRNeasy kit (Qiagen, 217004). RNA concentration was measured using a NanoDrop One/OneC Microvolume UV-Vis Spectrophotometer (ThermoFisher Scientific, ND-ONE-W). Reverse transcription was performed using SuperScript™ III Reverse Transcriptase (ThermoFisher Scientific, 18080-044) and 0.5 or 1 µg total RNA as a template with both oligo-dT (250ng) and random hexamers (100ng), unless stated otherwise. Quantitative-RT-PCR was performed by using a SYBR Green master mix (Applied Biosystems, Carlsbad, CA). Normalization of gene expression was performed by using the ΔΔCt method and statistical analysis by *t* testing. *TATA-box binding protein* (*TBP*) and *glyceraldehyde-3-phosphate dehydrogenase* (*GAPDH*) were used as reference genes for normalization. The list of DNA primers used in this study is provided in supplementary Table S7.

### Nucleo-cytoplasmic fractionation

Nucleo-cytoplasmic fractionation was performed using the PARIS kit (Ambion, ThermoFisher Scientific, AM1921) based on the manufacturer’s instructions. Briefly, HuCCT1 cells were trypsinized, pelleted and washed in PBS. Then, cells were gently resuspended in Cell Fractionation Buffer and incubated on ice for 10 min. Cell extracts were centrifuged at 500 × g at 4°C for 5 min and the cytoplasmic fraction was transferred to new tubes. The remaining nuclear pellet was washed once in Cell Fractionation Buffer and lysed in ice-cold Cell Disruption Buffer. RNA was isolated by adding a 2 × Lysis/Binding solution to each fraction, followed by addition of 100% ethanol and capture of the RNA by a filter cartridge. After a series of washes, the RNA was eluted in pre-heated Elution solution (PARIS kit) and stored at −80°C.

### Gene expression profiling

Total RNA was purified with an miRNeasy kit (Qiagen, 217004). Genome-wide expression profiling was performed using the low-input QuickAmp labeling kit and human SurePrint G3 8×60K pangenomic microarrays (Agilent Technologies, Santa Clara, USA) as previously described (Merdrignac *et al*, 2018). Differentially expressed genes were identified by a 2-sample univariate *t* test and a random variance model as previously described. Differentially regulated genes between the different experimental conditions were selected based on stringent criteria e.g. *P* value (*P*<0.001) and fold change (FC>2).

### *In vitro* RNA transcription

*In vitro* transcription of *LINC00313* or firefly luciferase (F-luc) mRNA was performed using the HiScribe™ T7 High Yield RNA Synthesis kit (New England Biolabs), according to the instructions by the manufacturer. Briefly, the pcDNA3.1-LINC00313 plasmid vector was linerized with single digestion, downstream of the insert site and the purified linearized plasmid was used as a template for *in vitro* transcription at 37°C for 2h. Residual DNA was degraded using RNase-free DNase I (New England Biolabs) and the *in vitro* synthesized RNA was purified using the RNA clean-up protocol from RNeasy Mini kit (Qiagen). The size and the integrity of the *in vitro* transcribed RNAs were verified using agarose gel electrophoresis.

### RNA labeling and RNA pull-down

The *in vitro* transcribed RNAs were labelled at the 3′ terminus with a desthiobiotinylated cytidine biphosphate nucleotide, using the Pierce RNA 3′ End Desthiobiotinylation kit (Pierce/Thermo Fisher Scientific, 20163), and were purified using the RNA cleanup protocol (QIAGEN). RNA pull-down assays were performed following the Pierce Magnetic RNA-Protein Pull-Down kit protocol (Pierce/Thermo Fisher Scientific, 20164). Biotinylated RNAs were pre-coupled to Nucleic Acid Compatible Streptavidin Magnetic Beads in RNA Capture Buffer (Pierce/Thermo Fisher Scientific) for 3h at 4°C. The RNA-beads complex was incubated with protein extracts from HuCCT1 cells, in the presence of Halt protease inhibitor cocktail (100X) (Thermo Fisher Scientific, 87786) and RNase inhibitor (Superase In, Ambion, Thermo Fisher Scientific, AM2694) for 1h at 4°C, with rotation. Next, the protein-RNA-beads complexes were washed and proteins were eluted in Elution Buffer (Pierce/Thermo Fisher Scientific, Stockholm, Sweden), boiled at 95°C for 5 min and subjected to SDS-PAGE, followed by immunoblotting for the detection of specific protein interactors. For the identification of *LINC00313*-interacting proteins the same protocol was applied followed by an unbiased proteomic analysis. In the latter case, the ribonucleoprotein complexes bound to beads were washed several times before being dissociated, using on-beads digestion and prepared for mass spectrometry analysis.

### Mass spectrometry

The analysis was performed by the Clinical Proteomics Mass Spectrometry facility (Karolinska Institutet, Karolinska University Hospital, Science for Life Laboratory, Stockholm, Sweden), as previously described (Papoutsoglou *et al*, 2019b, 2).

### RNA immunoprecipitation (RIP)

RIP was performed according to the Magna-RIP^TM^ RNA-binding protein immunoprecipitation kit (Millipore/Merck) as previously described (Papoutsoglou *et al*, 2019b).

### Immunoblotting

Total proteins were extracted using RIPA buffer (ThermoFisher Scientific, 89901) supplemented with protease and phosphatase inhibitor cocktails (ThermoFisher Scientific, 13393126). Cell lysates were briefly sonicated and cleared by centrifugation (10,000 g, 15 min). Protein concentration was determined, using the Pierce bicinchoninic acid (BCA) Protein Assay kit (ThermoFisher Scientific, 23227). Then, 4X NuPAGE LDS sample buffer (ThermoFisher Scientific, NP0007), supplemented with 10X NuPAGE reducing agent (ThermoFisher Scientific, NP0009) was added to the lysates, which were then boiled at 70°C for 10 min. Protein samples (20 or 40 µg) were loaded in NuPAGE™ Novex™ 4-12% Bis-Tris Protein Gels, 1.0 mm, 10-well (ThermoFisher Scientific, NP0321) and the resolved proteins were transferred to a nitrocellulose filter using the iBlot Dry Blotting System (Invitrogen, IB1001). Then, the filters were blocked in 5% BSA/Tris-buffered saline (TBS) containing 0.1% Tween-20 or 5% nonfat dry milk (Cell Signalling Technology, 9999S) in TBS-Tween-20 and incubated with primary antibody solutions (overnight, 4°C). Anti-rabbit (Cell Signalling Technology, #7074) or anti-mouse (Cell Signalling Technology, #7076) horseradish peroxidase-conjugated secondary antibodies were incubated with membranes for 1 h at room temperature and blots were developed, using enhanced chemiluminescence (ECL) assays (Cytiva,). Images were taken using an ImageQuant LAS 4000 imager (GE Healthcare). A list of the primary antibodies used in this study is provided in Supplementary Table S8.

### Immunofluorescence

Empty vector or LINC00313 overexpressing HuCCCT1 cells were seeded in 96-well plates and fixed in 4% PFA for 15 min at room temperature. After three washes in PBS, cells were blocked in 5% FBS/PBS, supplemented with 0.3% Triton X-100 for 1h at room temperature. Then, samples were incubated with primary antibodies (anti-β-catenin, anti-ACTL6A) in 1% BSA/PBS solution overnight at 4 °C. The next day, samples were washed three times in PBS and Dylight 488-conjugated goat anti-rabbit secondary antibody (Insight Biotechnology, 5230-0385) in 1% BSA/PBS solution was added to the cells together with Hoechst stain for nuclear staining. Images were acquired with an EVOS M5000 Imaging System (ThermoFisher Scientific, AMF5000).

### Luciferase assays

Stable TGFβ sensor HuCCT1 cells expressing a TGFβ-responsive luciferase reporter construct (HuCCT1-Lenti-SMAD), have previously been described (Merdrignac *et al*, 2018). Luciferase reporter assays were performed using the Luciferase Assay System (Promega, E1501), according to the manufacturer’s instructions. Briefly, 20,000 HuCCT1-Lenti-SMAD or HuCCT1-Lenti-NC (TGFβ-unresponsive, negative control) cells were seeded in 96-well plates and treated with TGFβ and with or without MEK, JNK, p38 and PI3K inhibitors for 16 h. For TCF/LEF-luciferase reporter assays, HuCCT1 cells stably expressing pGreenFire 2.0 TCF/LEF reporter virus (pGF2-TCF/LEF-rFluc-T2A-GFP-mPGK-Puro) were silenced, over-expressed or treated with inhibitors or TGFβ as described in the figure legends. Luciferase activity was normalized to the number of viable cells measured by a PrestoBlue cell viability reagent (ThermoFisher Scientific, A13261). When using TOPFlash or FOPFlash for TCF/LEF-luciferase reporter assays, cells were transiently transfected with each one of TOPFlash or FOPFlash together with pGL4.74 [hRluc/TK] Vector, expressing Renilla luciferase. Firefly luciferase activity was normalized to Renilla luciferase activity using the Dual-Luciferase Reporter Assay System (Promega, E1910). Luciferase activity measurements were performed using a TECAN Spark multimode microplate reader. Each experiment was performed in biological quadruplicates.

### Functional tests

Cell viability was measured using a PrestoBlue reagent (ThermoFisher Scientific, A13261) according to the manufacturer’s instructions. Cell migration was evaluated, with wound healing assays at different time points (0, 8, 24 hours) using silicone inserts with a defined cell-free gap (Ibidi, 80366). After the removal of the insert, cells were grown in 0.1% FBS/RPMI, supplemented with 10 µg/ml mitomycin C (Sigma-Aldrich, M4287), in order to stop their proliferation. Wound closure was evaluated using the ImageJ software. For colony formation assays, 800 HuCCT1 cells were seeded in 6-well plates and cultured for 11 days. Colonies were washed in PBS twice and fixed in 4% paraformaldehyde (PFA) for 15 min. Then, colonies were stained with 0.1% crystal violet and counted manually. Each experiment was performed in biological triplicates.

### *In vivo* xenografts

HuCCT1 cells over-expressing empty vector (pcDNA3.1) or LINC00313 (pcLINC00313) were infected with GL261-Luc (CMV-Firefly luciferase lentivirus (Neo), PLV-10064-50, Cellomics Technology, USA). 4,000,000 cells (100µL in PBS + 100µL matrigel) were implanted on the flanks of immunodeficient NSG (*NOD Cg-Prkdcscid Il2rgtm1Wjl/SzJ*) 8-week-old male mice (Charles River, USA). All animal procedures met the European Community Directive guidelines (Agreement B35-238-40, Biosit Rennes, France; DIR #7163) and were approved by the local ethics committee and ensuring the breeding and the daily monitoring of the animals in the best conditions of wellbeing according to the law and the 3R rule (Reduce-Refine-Replace). Tumor growth was evaluated by a direct measurement of tumor size using caliper. Mice were sacrificed 77 days after implantation and tumor, liver and lung tissues were isolated and subjected to molecular analysis.

### RNA sequencing of mice xenograft tumors

Xenograft tumors were cut into smaller pieces in a Petri dish placed on ice and transferred into 2mL tubes containing 2.8mm ceramic beads (P000911-LYSK0-A, Bertin Corp) filled with 1 ml Qiazol. Homogenization of the tumors was performed using the Precellys 24 tissue homogenizer (P000669-PR240-A, Bertin Instruments) at 5,500-rpm speed, 3 cycles of 20s each with a pause of 20s, between the cycles. Then, the homogenate was transferred to new tubes and subjected to RNA extraction using the RNeasy Lipid Tissue Mini Kit (74804, Qiagen), according to the instructions by the manufacturer. RNA-seq was performed by BGI Genomics. The RNA-seq data have been deposited to ArrayExpress with accession number E-MTAB-12030.

### Integrative analysis

We used the intrahepatic cholangiocarcinoma RNA-Seq data from Dong et al. (Dong *et al*, 2022) publicly available in the NODE database under the accession number OEP001105 (https://www.biosino.org/node/project/detail/OEP001105). Raw counts were normalised with the DESeq2 and scaled. We then performed a Principal Component Analysis (PCA) on the OEP001105 normalised data with samples from our study as supplementary individuals (datasets were merged based on the gene symbols of the signature). We used an agglomerative hierarchical clustering on the PCA components (Hierarchical Clustering on Principle Components) to divide the samples from the OEP001105 dataset into two groups: CTRL (pcDNA3) or LINC (pcLINC00313). Overall survival (OS) was estimated using the Kaplan-Meier method. Comparison between survival groups was performed using a log-rank test for binary variables. All the analyses were conducted with the following R packages: DESeq2, FactoMineR and survival.

### Statistical analysis

Comparisons were performed using a two-tailed paired Student’s t test (*P < 0.05, **P < 0.01, ***P < 0.001). The results are shown as mean ± SD from 3 independent experiments.

## Acknowledgements

We thank Dr Yutaro Tsubakihara and Prof. Aristidis Moustakas (Uppsala University, Sweden) for providing the pcDNA3.1/Hygro (+) and the pGL4.74 TK-Renilla luciferase reporter expression vectors. We thank Dr Laura Fouassier for providing CCA cell lines and for helpful discussion. Gaëlle Angenard, Raffaële Leroux, Léa Chailloux and Matthis Desoteux are acknowledged for their technical help. The authors are supported by Inserm, Université de Rennes 1, Région Bretagne, Ministère de l’Enseignement Supérieur de la Recherche et de l’Innovation, Ligue Nationale Contre le Cancer, Ligue Contre le Cancer (CD22, CD35, CD44, CD85), Fondation ARC, INCa and ITMO Cancer AVIESAN (Alliance Nationale pour les Sciences de la Vie et de la Santé) dans le cadre du Plan cancer (Non-coding RNA in cancerology: fundamental to translational). This work was also supported by a grant from the French Ministry of Health and the French National Cancer Institute, PRT-K20-136, CHU Rennes, CLCC Eugène Marquis, Rennes. JMB is supported by Spanish Carlos III Health Institute (ISCIII) [(FIS PI18/01075, PI21/00922, and Miguel Servet Programme CPII19/00008) cofinanced by “Fondo Europeo de Desarrollo Regional” (FEDER)] and CIBERehd (ISCIII); La Caixa Scientific Foundation (HR17-00601); AMMF-The Cholangiocarcinoma Charity (EU/2019/AMMFt/001); PSC Partners US and PSC Supports UK (06119JB); European Union’s Horizon 2020 Research and Innovation Program [grant number 825510, ESCALON].

## Author contributions

CC conceived the project, provided funding and supervision. PP and CC designed the experiments. PP, CL, and RP acquired the data. PP, CL, RP, ALH and MA analysed the data. PP, CL, RP, ALH, MA and CC interpreted the data. JMB and DG contributed resources and editing. PP drafted the article. All authors critically revised the article.

## Data availability statement

*In vivo* and *in vitro* RNA-seq data, as well as microarray and ATAC-seq data are available at ArrayExpress under the accession number E-MTAB-12030. The ATAC-seq peaks can be visualized in the UCSC Genome Browser following the URL: https://genome.ucsc.edu/s/ppapouts/ATAC%2Dseq%20HuCCT1%20LINC00313%20overexpression.

**Expanded view** for this article is available online.

## SUPPLEMENTAL INFORMATION

### Supplementary Figures

**Supplementary Fig. S1.**
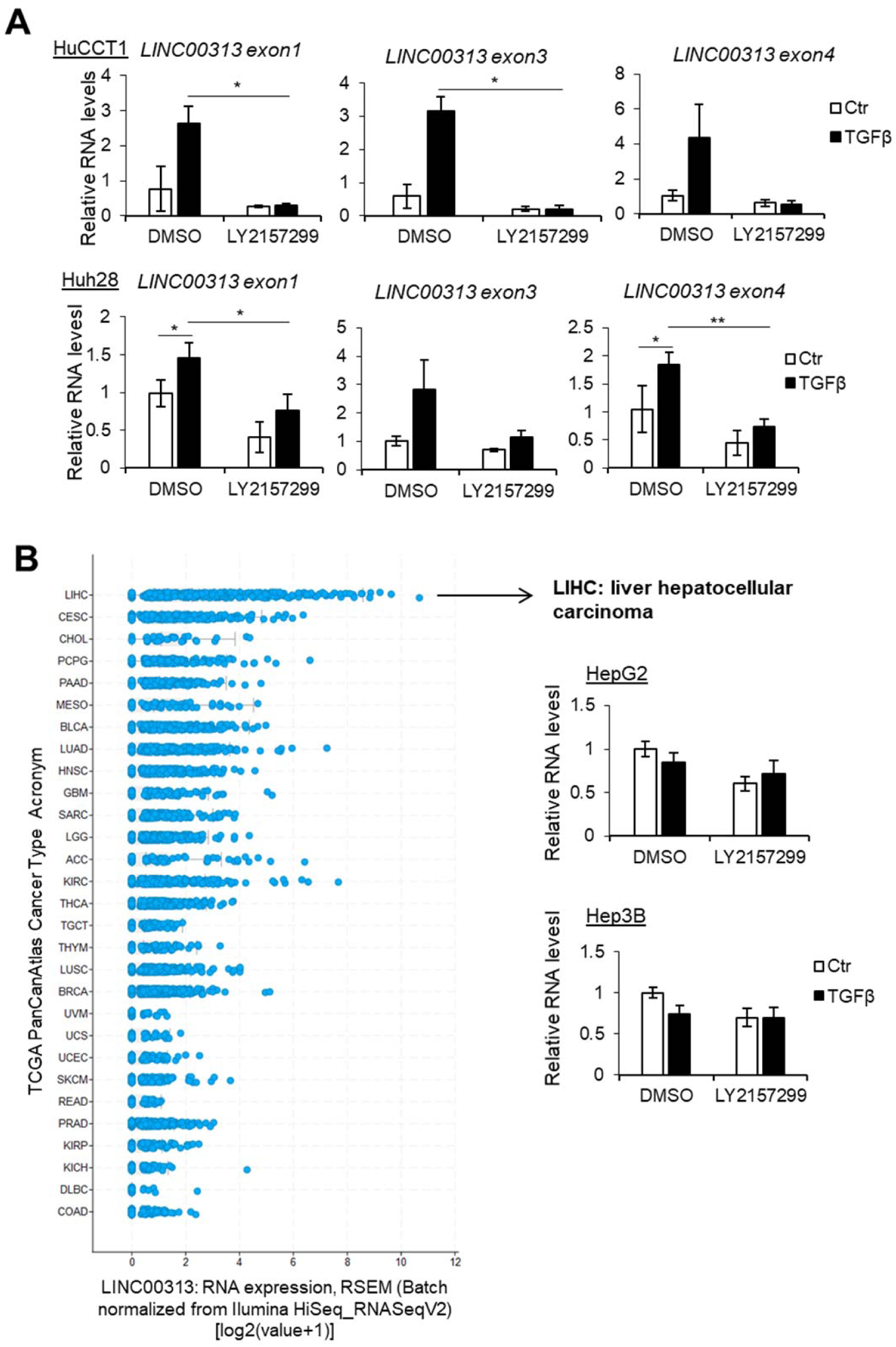
TGFβ-mediated *LINC00313* up-regulation is CCA-specific. A) Real-time qPCR for detection of LINC00313 specific exonic regions in HuCCT1 or Huh28 cells treated with the TβRI inhibitor LY2157299 or DMSO, and stimulated with TGFβ1 or BSA/HCl for 16h. B) LINC00313 expression levels (normal logarithmic scale) among every cancer type of the PanCancer Atlas of TCGA, sorted from the highest (top) to the lowest (bottom) expression and real-time qPCR to detect LINC00313 expression in HCC cell lines HepG2 and Hep3B treated as in panel A.

**Supplementary Fig. S2.**
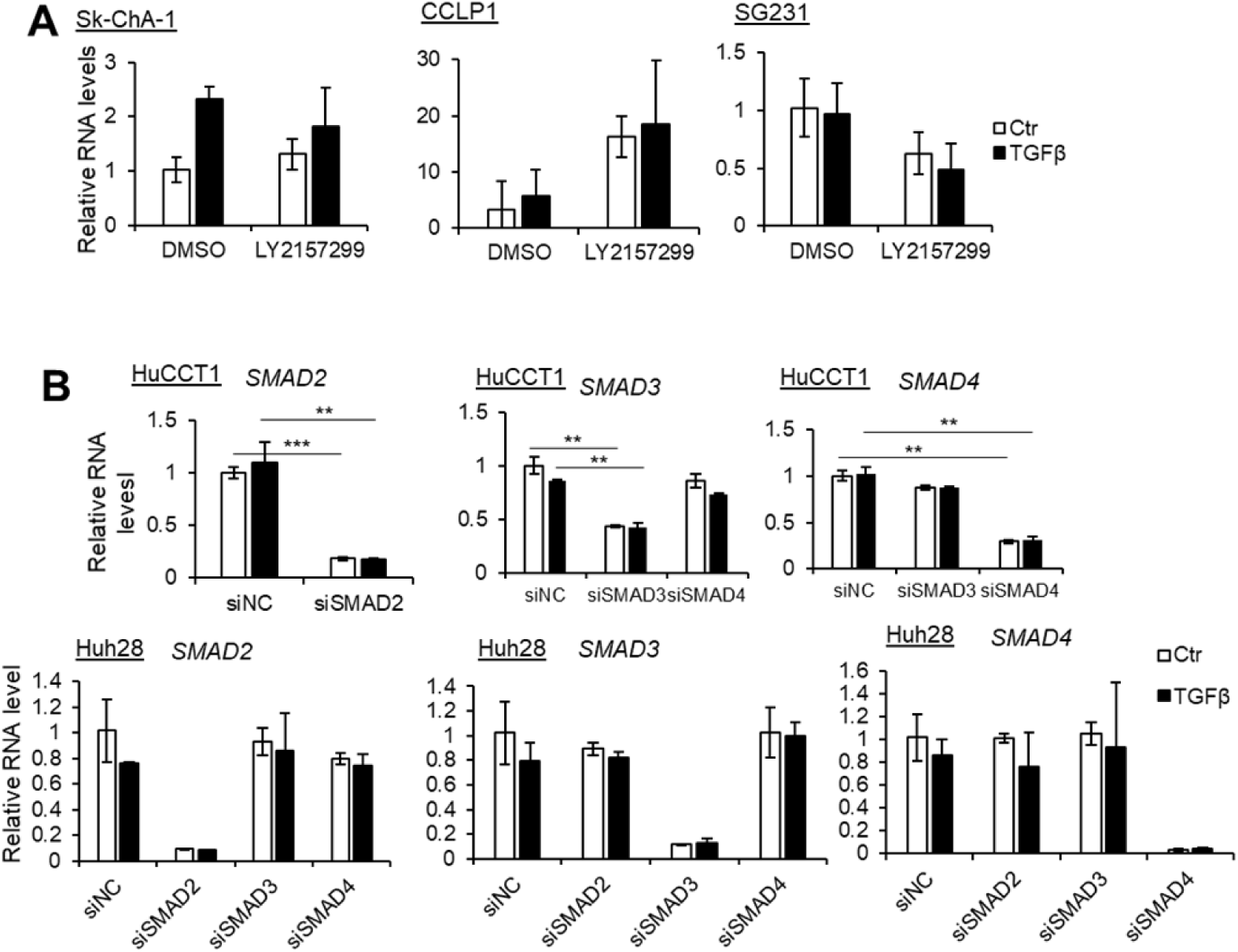
*LINC00313* is induced by TGFβRI-SMAD signalling. A) Real-time qPCR for detection of *LINC00313* in eCCA cell line Sk-ChA-1 and in iCCA cell lines CCLP1, SG231 treated with the TβRI inhibitor LY2157299 or DMSO, and stimulated with TGFβ1 or BSA/HCl for 16h. B) Real-time qPCR to measure *SMAD2*, *SMAD3* and *SMAD4* mRNA levels in HuCCT1 and Huh28 cell lines transiently transfected with siRNAs targeting *SMAD2*, *SMAD3* or *SMAD4* or a non-targeting siRNA (siNC) and stimulated with TGFβ1 or BSA/HCl for 16h.

**Supplementary Fig. S3.**
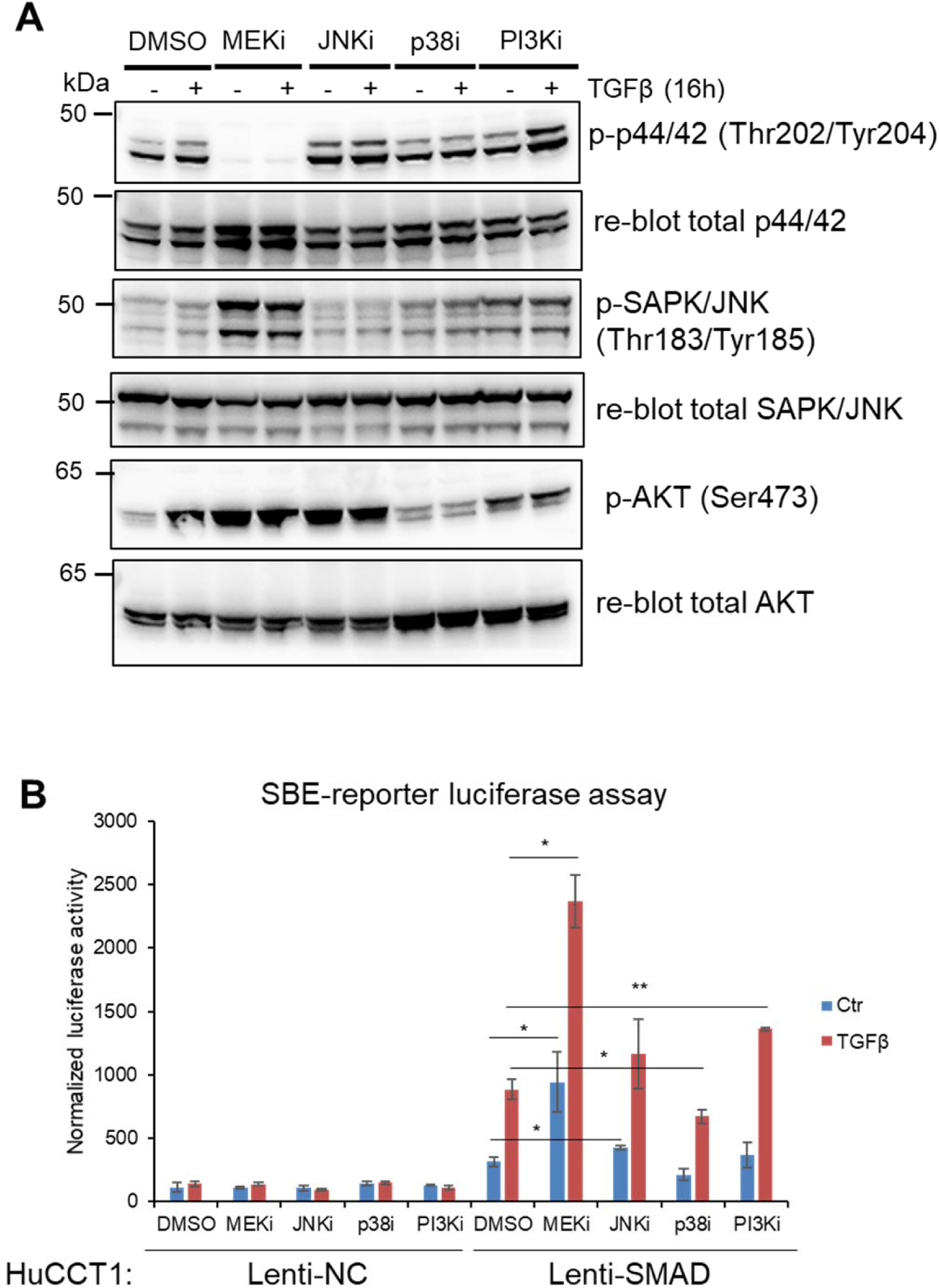
Inhibition of MAPKs in HuCCT1 cells. A) Immunoblotting for detection of phosphorylated p44/42 (Thr202/Tyr204), phosphorylated p-SAPK/JNK (Thr183/Tyr185) and phosphorylated AKT (Ser473) in HuCCT1 cells treated with MEK, p38, JNK or PI3K inhibitors, with or without TGFβ1 stimulation for 16h. DMSO was used as a vehicle treatment. B) SBE-luciferase reporter assay in HuCCT1 cells treated with MEK, p38, JNK or PI3K inhibitors, with or without TGFβ1 stimulation for 16h. DMSO was used as a vehicle treatment.

**Supplementary Fig. S4.**
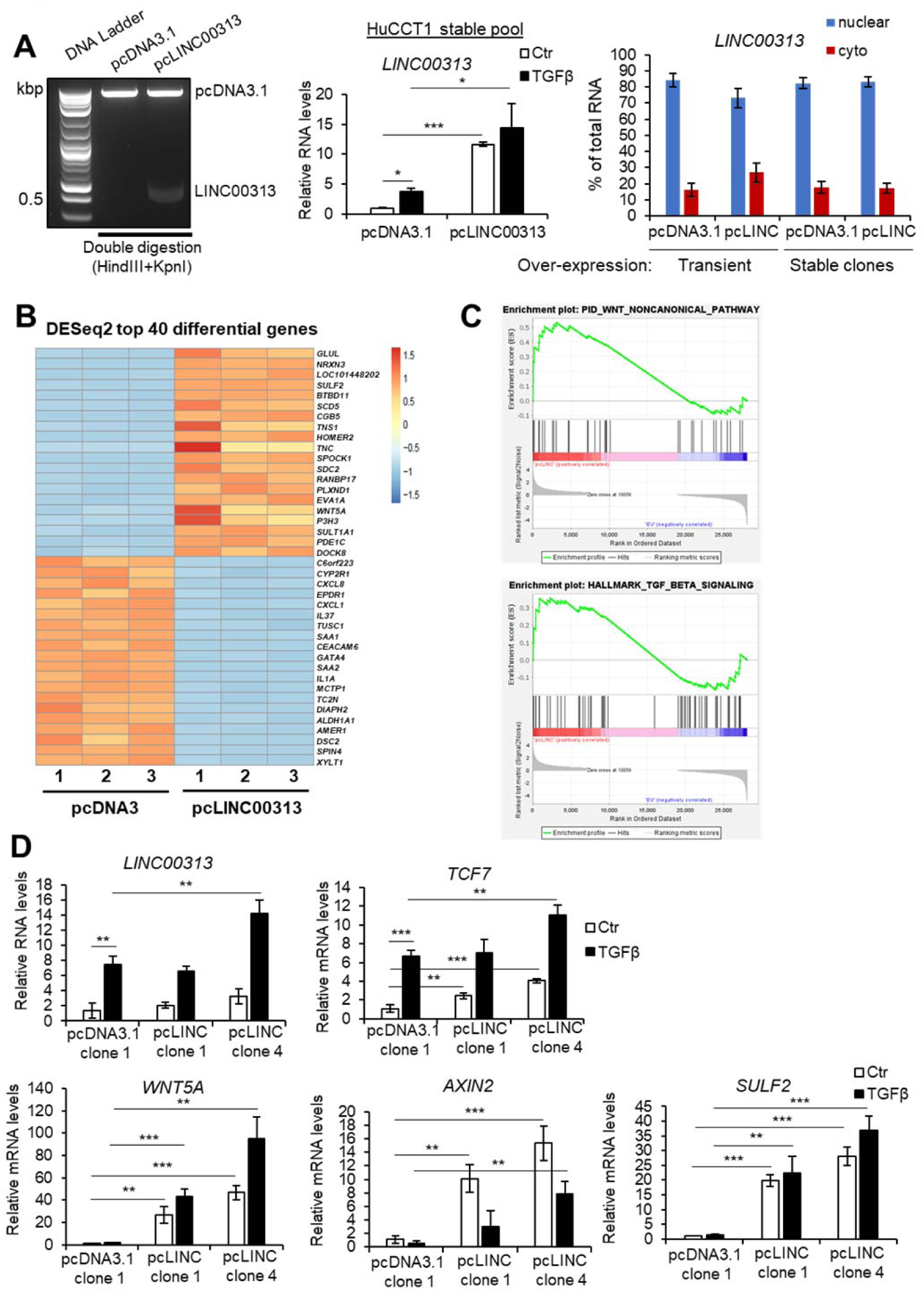
LINC00313 gain of function up-regulates Wnt-related genes. A) Efficient ligation of *LINC00313* insert to pcDNA3.1 plasmid vector was verified by double digestion with restriction enzymes, followed by agarose gel electrophoresis. Molecular size (in kbp) marker ladder is shown in the first lane. Real-time qPCR analysis of *LINC00313* expression in HuCCT1 cells stably over-expressing pcDNA3.1 or pcLINC00313 expression vectors. Subcellular fractionation followed by RT-qPCR analysis of *LINC00313* RNA levels in nuclear and cytoplasmic fractions of HuCCT1 cells transiently over-expressing an empty vector (pcDNA3.1) or *LINC00313* and HuCCT1 clones, stably over-expressing pcDNA3.1 or LINC00313. B) Heatmap presenting the top 40 differentially regulated genes in HuCCT1 pcLINC00313 versus pcDNA3-expressing cells (shrunken log2FC>1, adjusted p-value<0.1). Biological triplicates (1-3) were used per condition. C) GSEA analysis of differentially expressed genes in pcLINC00313 versus pcDNA3 HuCCT1 cells. D) Real-time qPCR analysis of *LINC00313*, *TCF7*, *WNT5A*, *AXIN2*, and *SULF2* expression in a monoclonal HuCCT1 cell line stably expressing pcDNA3.1 and in two monoclonal HuCCT1 cell lines stably expressing pcLINC00313, in the presence or not of TGFβ1 for 16h.

**Supplementary Fig. S5.**
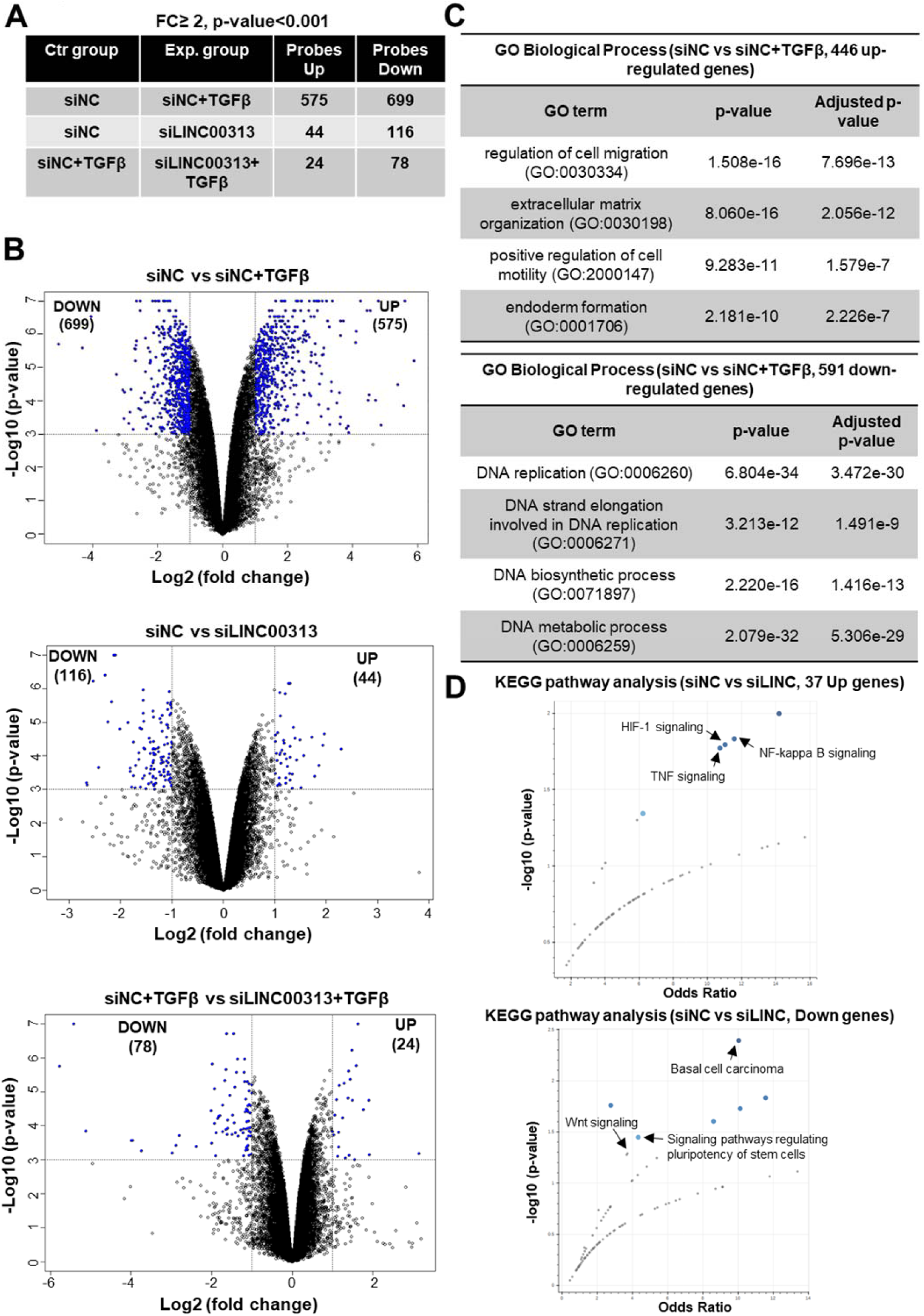
Gene expression profiling after *LINC00313* loss of function. A) Number of differentially expressed genes in HuCCT1 cells transiently transfected with an individual siRNA targeting *LINC00313* or a non-targeting siRNA (siNC) and stimulated with TGFβ1 or not for 16h. B) Volcano plots depicting the differentially up- or down-regulated genes between different experimental conditions C) Gene ontology analysis of the 446 up-regulated and 591 down-regulated genes, in response to TGFβ1 stimulation. The top four statistically significant GO terms, related to biological process, are shown in the table, together with p-value and adjusted p-value for each GO term. D) KEGG pathway analysis of differentially up- or down-regulated genes in response to *LINC00313* silencing in HuCCT1 cells.

**Supplementary Fig. S6.**
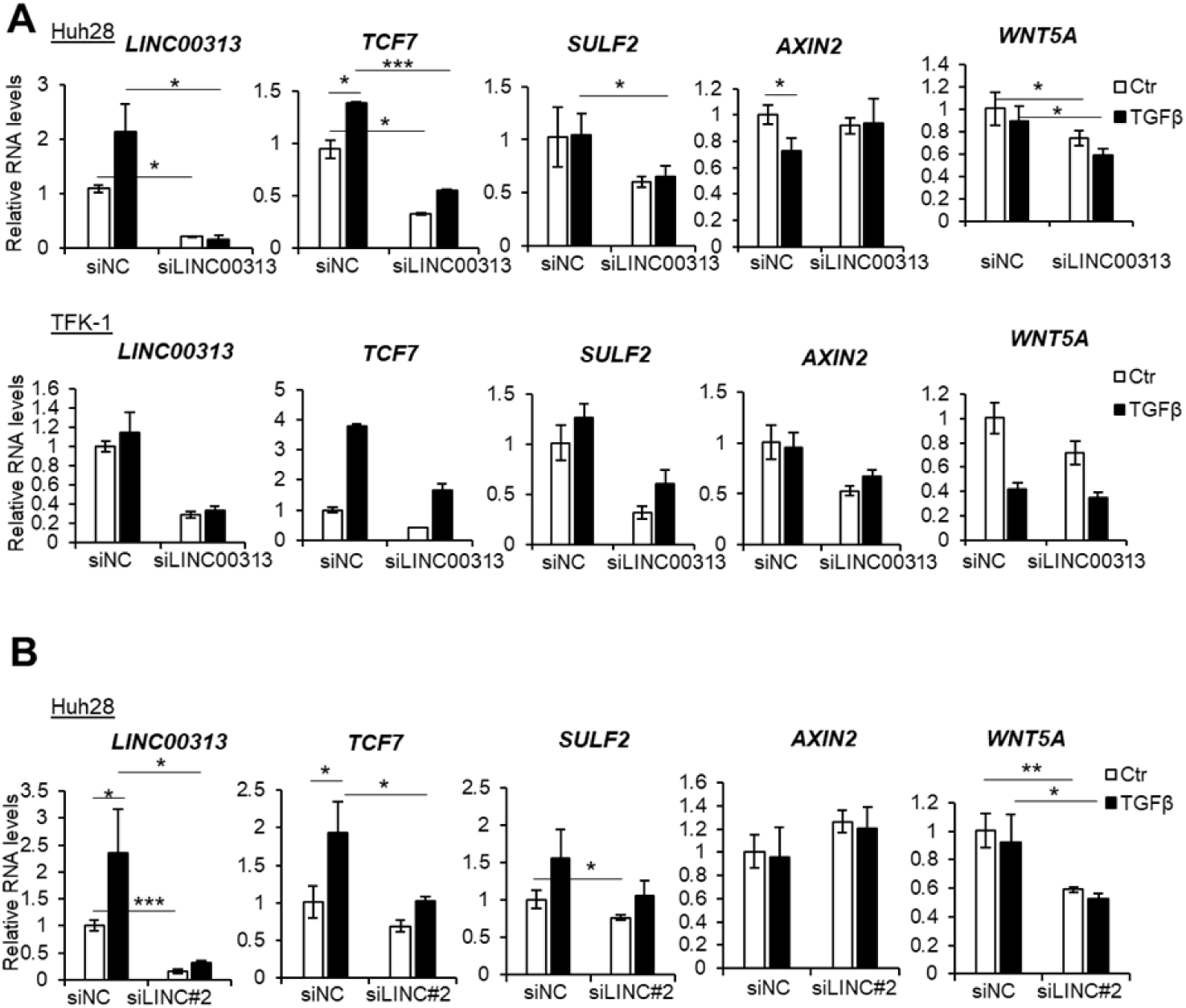
Silencing *LINC00313* in CCA cell lines regulates target gene expression. A) Real-time qPCR analysis of *LINC00313*, *TCF7*, *SULF2*, *AXIN2* and *WNT5A* expression in Huh28 and TFK-1 cells, transiently transfected with an individual siRNA targeting *LINC00313* (siLINC00313) or a non-targeting siRNA (siNC) and stimulated or not with TGFβ1 for 16h. B) Real-time qPCR analysis of *LINC00313*, *TCF7*, *SULF2*, *AXIN2* and *WNT5A* expression in Huh28 cells, transiently transfected with a second individual siRNA targeting *LINC00313* (siLINC#2) or a non-targeting siRNA (siNC) and stimulated or not with TGFβ1 for 16h.

**Supplementary Fig. S7.**
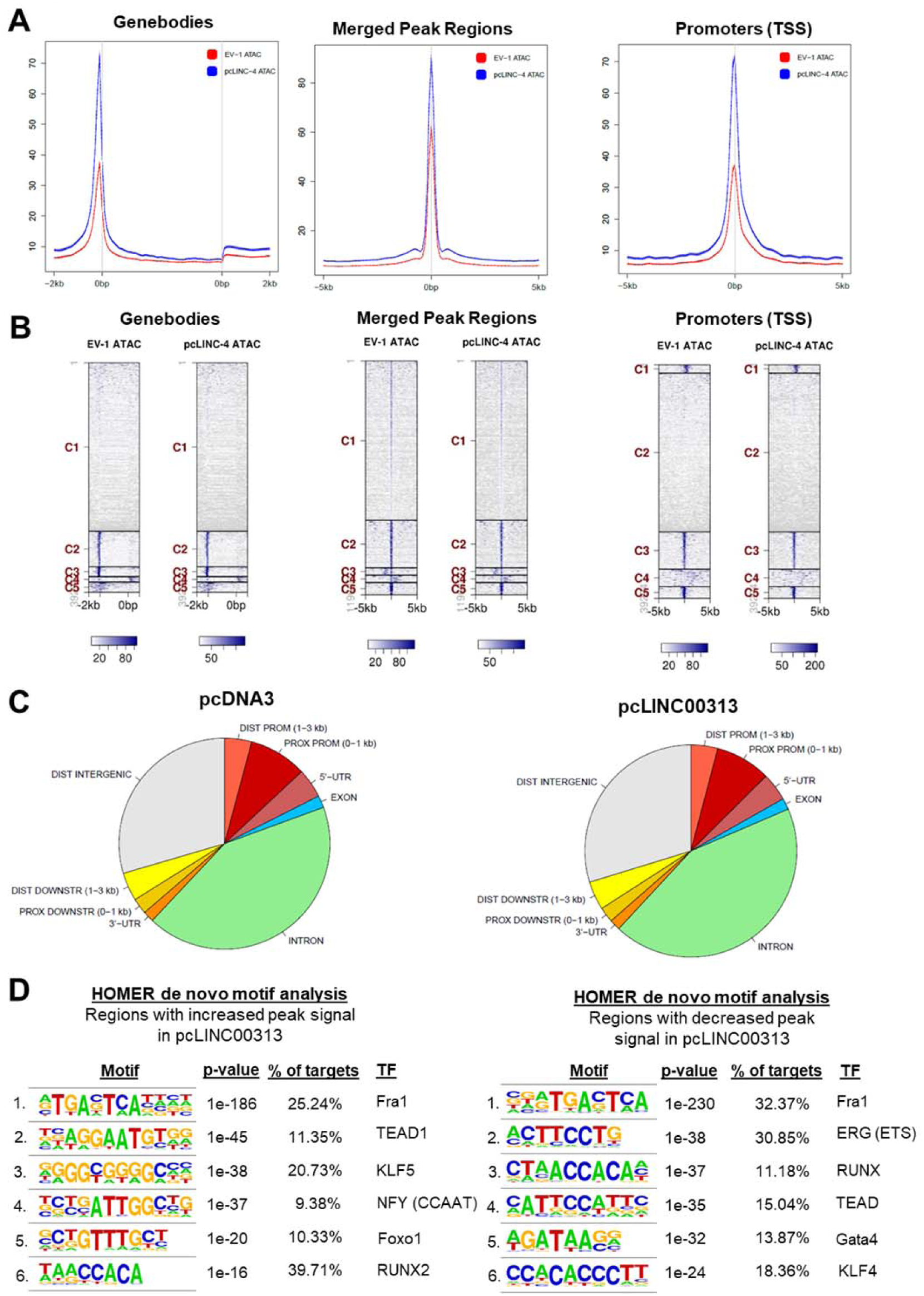
*LINC00313* does not cause global alterations on chromatin accessibility. A-B) Tag distributions across target regions such as Merged Regions (= all peak regions; +/- 5 kb), transcription start sites (TSS; +/- 5 kb) or gene bodies (with 2 kb flanking regions) are presented as average plots (A) and as heatmaps (B). C) Piecharts displaying the location of ATAC-seq peaks relative to genomic annotations. Several peaks are assigned to more than one feature. D) HOMER-based analysis of transcription factor binding motifs in chromatin regions with increased or decreased peak signal after *LINC00313* over-expression. The top 6 most significant consensus motifs, together with the percentage of target regions that are assigned to, are depicted.

**Supplementary Fig. S8.**
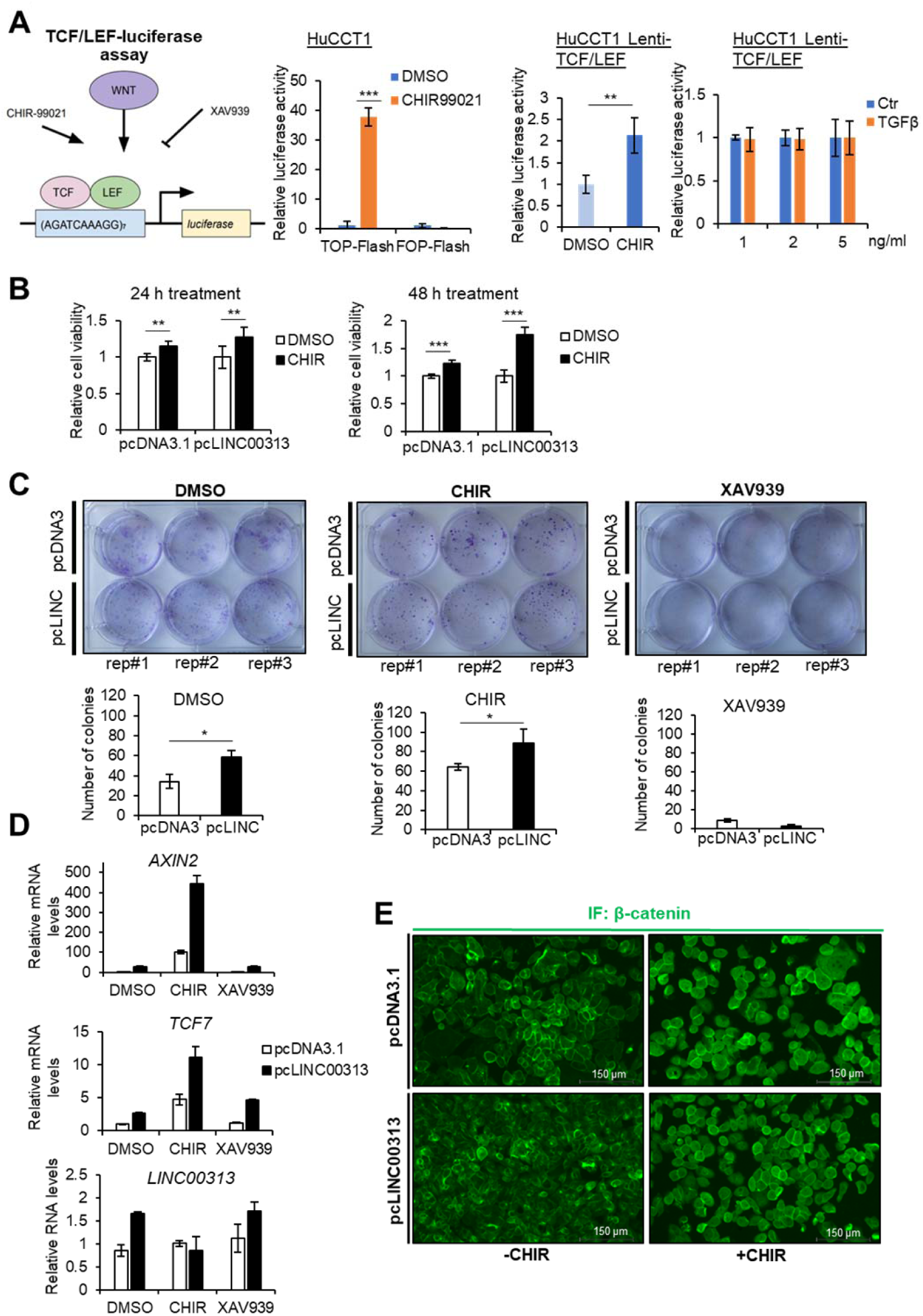
Chemically-induced Wnt/β-catenin signalling impacts CCA physiological processes. A) Schematic representation of the TCF/LEF-luciferase reporter assay. Activators (Wnt ligands, CHIR) and inhibitors (XAV939) of TCF/LEF-dependent responses are also shown. TCF/LEF-luciferase reporter assay in HuCCT1 cells transiently transfected with TOP-Flash or FOP-Flash luciferase expression vectors and treated with CHIR99021 or DMSO for 24h. Cells were co-transfected with a Renilla luciferase expression vector for normalization of the firefly luciferase activity. TCF/LEF-luciferase reporter assay in HuCCT1 cells stably expressing the pGreenFire 2.0 TCF/LEF reporter construct and treated with CHIR99021 or DMSO for 24h. TCF/LEF-luciferase reporter assay in HuCCT1 cells stably expressing the pGreenFire 2.0 TCF/LEF reporter construct and treated with the indicated concentrations of TGFβ1 or BSA/HCl (Ctr) for 16h. B) Cell viability assays in control or *LINC00313* over-expressing HuCCT1 cells treated with CHIR99021 or DMSO for 24h. Six biological replicates were used per condition. C) Colony formation assay in control or *LINC00313* over-expressing HuCCT1 cells, treated with CHIR99021 or XAV939 or DMSO for 24h. Quantification of the number of colonies for each condition is also shown. D) Real-time qPCR analysis of *AXIN2*, *TCF7* and *LINC00313* expression in control or *LINC00313* over-expressing HuCCT1 cells, treated with CHIR99021, XAV939, or DMSO for 24h. E) Immunofluorescence to detect β-catenin subcellular localization in control or *LINC00313* over-expressing HuCCT1 cells, treated or not with CHIR99021 or DMSO for 24h.

**Supplementary Fig. S9.**
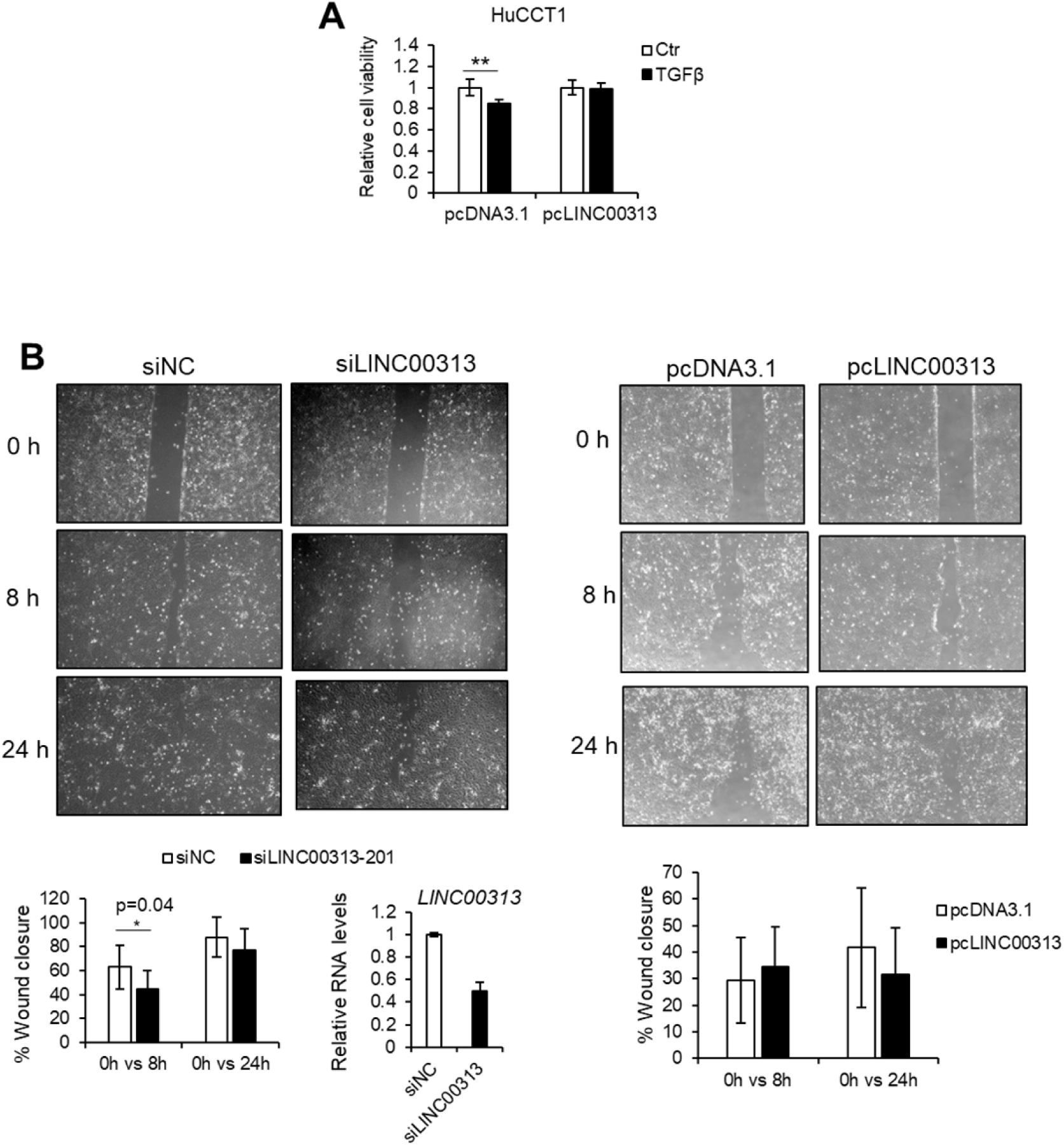
*LINC00313* affects HuCCT1 cell viability but not migration. A) Cell viability assay in control or *LINC00313* over-expressing HuCCT1 cells stimulated or not with TGFβ1 for 16h. B) Wound healing assay in HuCCT1 cells, transiently transfected with an individual siRNA targeting *LINC00313* or a non-targeting siRNA (siNC) and in pcDNA3 and pcLINC00313 over-expressing cells. Quantification of migration is shown as percentage of wound closure after eight and 24h since the scratch was performed. Real-time qPCR analysis of *LINC00313* to quantify the efficiency of *LINC00313* silencing is also shown.

**Supplementary Fig. S10.**
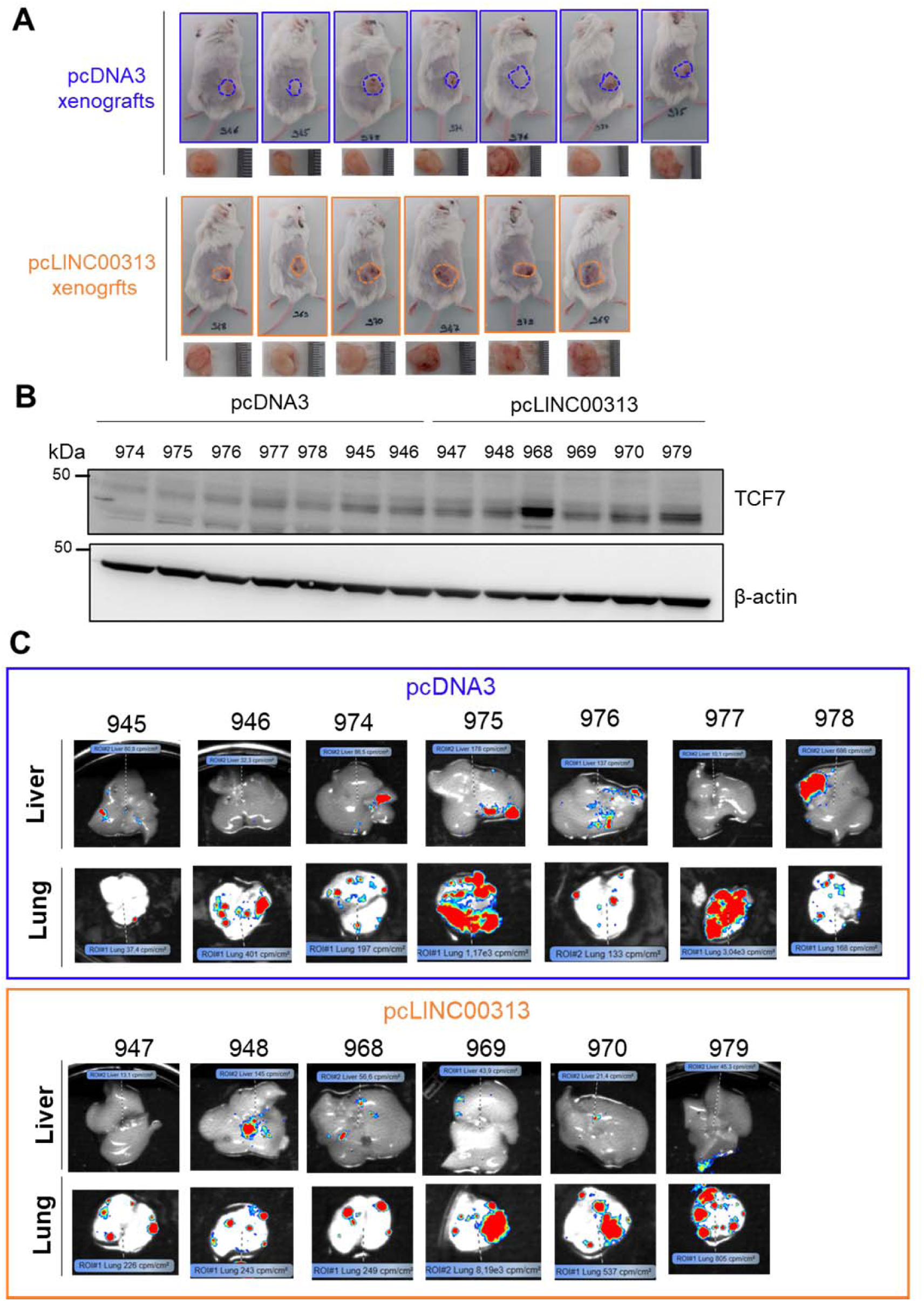
Impact of *LINC00313* on tumor growth and metastasis in a xenograft CCA mouse model. A) Picture of the mice and the corresponding resected tumours. B) Immunoblotting for detection of TCF7 and β-actin protein levels using total protein extracts from the resected tumours as described in panel A. C) Pictures showing the presence of metastases in resected livers and lungs of the same mice. Bioluminescent signal, derived from HuCCT1 cells expressing luciferase, indicates the sites of metastasis.

**Supplementary Fig. S11.**
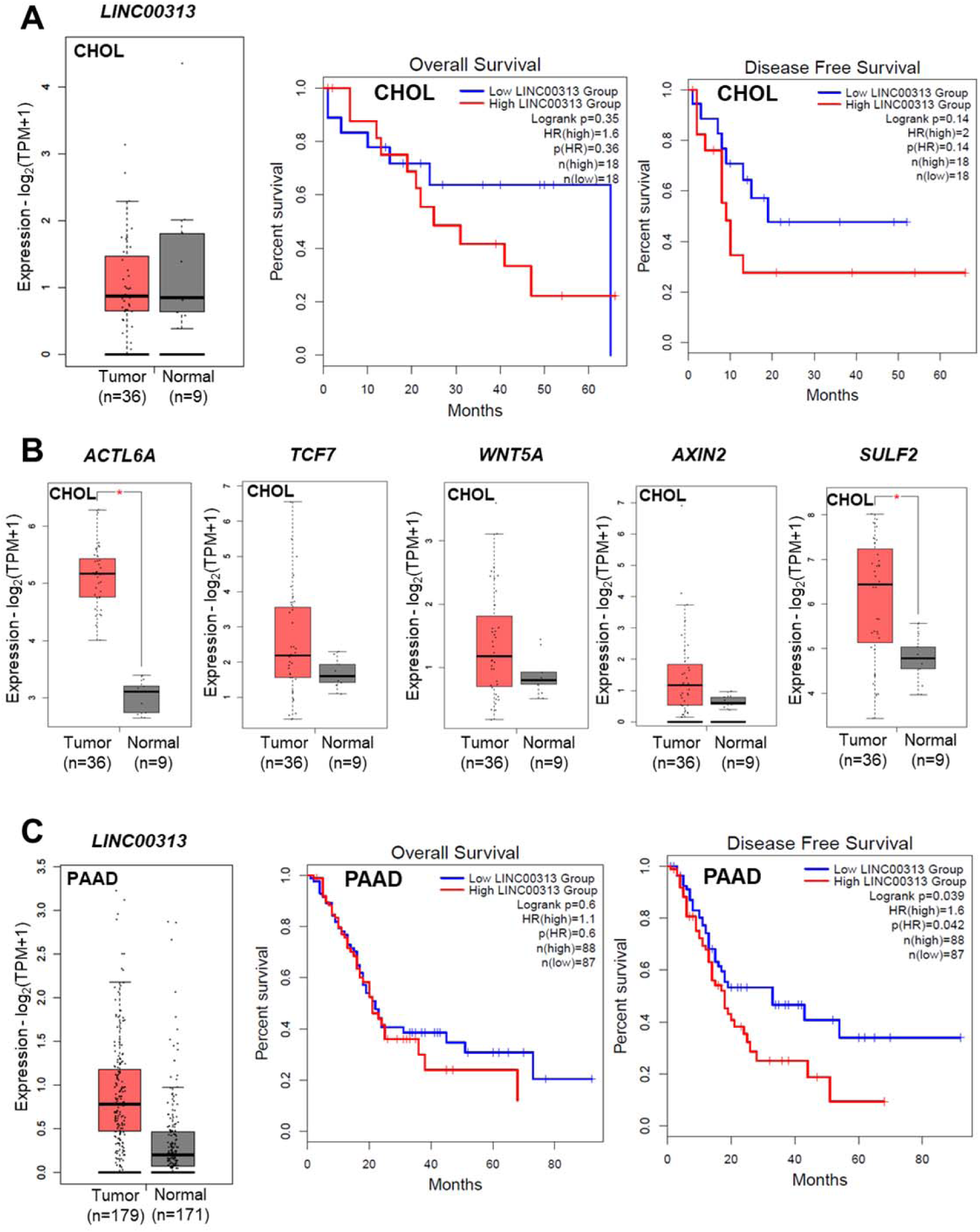
Clinical relevance of *LINC00313* in human CCA and pancreatic adenocarcinoma. A) Boxplot of *LINC00313* expression in TPM and Kaplan-Meier survival curves depicting overall and disease free survival in CHOL patients grouped based on *LINC00313* median expression. B) Boxplots of *ACTL6A*, *TCF7*, *WNT5A*, *AXIN2* and *SULF2* transcripts per million (TPMs) in cholangiocarcinoma (CHOL) tumors and their paired normal tissues. C) Boxplot of *LINC00313* expression in pancreatic adenocarcinoma (PAAD) tumors and their paired normal tissues and Kaplan-Meier survival curves depicting overall and disease free survival in PAAD patients grouped based on *LINC00313* median expression. TCGA and GTEx gene expression data and survival curves derived from GEPIA2. *p < 0.01.

